# The DNA helicase FANCJ (BRIP1) functions in Double Strand Break repair processing, but not crossover formation during Prophase I of meiosis in male mice

**DOI:** 10.1101/2023.10.06.561296

**Authors:** Tegan S. Horan, Carolline F. R. Ascenção, Christopher A. Mellor, Meng Wang, Marcus B. Smolka, Paula E. Cohen

## Abstract

During meiotic prophase I, recombination between homologous parental chromosomes is initiated by the formation of hundreds of programmed double-strand breaks (DSBs), each of which must be repaired with absolute fidelity to ensure genome stability of the germline. One outcome of these DSB events is the formation of Crossovers (COs), the sites of physical DNA exchange between homologs that are critical to ensure the correct segregation of parental chromosomes. However, COs account for only a small (∼10%) proportion of all DSB repair events; the remaining 90% are repaired as non-crossovers (NCOs), most by synthesis dependent strand annealing. Virtually all COs are formed by coordinated efforts of the MSH4/MSH5 and MLH1/MLH3 heterodimers. The number and positioning of COs is exquisitely controlled via mechanisms that remain poorly understood, but which undoubtedly require the coordinated action of multiple repair pathways downstream of the initiating DSB. In a previous report we found evidence suggesting that the DNA helicase and Fanconi Anemia repair protein, FANCJ (BRIP1/BACH1), functions to regulate meiotic recombination in mouse. A gene-trap disruption of *Fancj* showed an elevated number of MLH1 foci and COs. FANCJ is known to interact with numerous DNA repair proteins in somatic cell repair contexts, including MLH1, BLM, BRCA1, and TOPBP1, and we hypothesized that FANCJ regulates CO formation through a direct interaction with MLH1 to suppress the major CO pathway. To further elucidate the function of FANCJ in meiosis, we produced three new *Fancj* mutant mouse lines via CRISPR/Cas9 gene editing: a full-gene deletion, a mutant line lacking the MLH1 interaction site and the N-terminal region of the Helicase domain, and a C-terminal 6xHIS-HA dual-tagged allele of *Fancj.* We also generated an antibody against the C-terminus of the mouse FANCJ protein. Surprisingly, while Fanconi-like phenotypes are observed within the somatic cell lineages of the full deletion *Fancj* line, none of the *Fancj* mutants show any change in either MLH1 focus counts during pachynema or total CO number at diakinesis of prophase I of meiosis. We find evidence that FANCJ and MLH1 do not interact in meiosis; further, FANCJ does not co-localize with MSH4, MLH1, or MLH3 during late prophase I. Instead, FANCJ forms discrete foci along the chromosome cores beginning in early meiotic prophase I, occasionally co-localizing with MSH4, and then becomes densely localized on unsynapsed chromosome axes in late zygonema and to the XY chromosomes in early pachynema. Strikingly, this localization strongly overlaps with BRCA1 and TOPBP1. *Fancj* mutants also exhibit a subtle persistence of DSBs in pachynema. Collectively, these data suggest a role for FANCJ in early DSB repair events, and possibly in the formation of NCOs, but they rule out a role for FANCJ in MLH1-mediated CO events. Thus, the role of FANCJ in meiotic cells involves different pathways and different interactors to those described in somatic cell lineages.

## Introduction

Meiosis is the specialized cell division that produces haploid gametes in sexually reproducing organisms. At the onset of meiotic prophase I, SPO11 and its accessory proteins induce hundreds of programmed double strand breaks (DSBs) throughout the genome^1–3^. Although DSBs are highly toxic forms of DNA damage when induced spontaneously, they are a crucial first step to crossover (CO) formation during meiosis. Meiotic COs, sites of physical DNA exchange between homologous chromosomes, play an essential role in ensuring correct chromosome segregation and thus preventing aneuploidy. It stands to reason, therefore, that the distribution and frequency of COs across the genome are tightly regulated during meiotic prophase I, with only 10% of the ∼250 initiating DSBs being resolved as COs in the laboratory mouse. At the same time, it is crucial that all DSBs are repaired appropriately prior to exit from prophase I of meiosis. Following their induction, all DSB ends undergo resection, producing a 3’ single-stranded DNA tail on which the RecA homologs RAD51 and DMC1 form a nucleoprotein filament^4,5^. RAD51/DMC1 promote invasion of the 3’ strand into the homologous chromosome and the subsequent formation of a D-loop structure^6–8^, which can either be disassembled to generate non-crossovers (NCOs) through synthesis dependent strand annealing (SDSA) or produce joint molecule (JM) intermediates necessary for COs^9–11^. Of the SPO11-induced DSBs that form COs, most (90-95%) are resolved via the class I CO pathway, which utilizes members of the DNA mismatch repair (MMR) protein family: MutSγ (MutS homologs, MSH4/MSH5) first bind to and stabilize JMs before MutLγ (MutL homologs, MLH1/MLH3) catalyze the resolution of the JM to form a CO^12–15^. The remaining 5-10% of COs are catalyzed by structure-selective nucleases (SSNs), e.g., the class II COs formed by MUS81-EME1 which are thought to act on atypical JMs^16,17^. The regulatory process ensuring how the correct number and distribution of COs is ensured from the hundreds of DSBs, and how the two CO pathways are coordinated as part of this regulatory process, remain poorly understood.

We previously reported that the DNA helicase FANCJ (BRCA1 Interacting Protein 1 [BRIP1] or BRCA1-Associated C-Terminal Helicase [BACH1]) has a role in the regulation of meiotic COs^18^. Specifically, disruption of *Fancj* by gene-trap (*Fancj^GT/GT^*) caused an increase of MLH1 foci in pachytene spermatocytes and an overall increase in total COs formed. FANCJ is a member of the Fanconi Anemia (FA) network, a pathway consisting at least 22 genes. The FA network canonically catalyzes the repair of interstrand crosslinks (ICLs) during DNA replication, but work over the last decade has shown that the FA network additionally serves as a hub that bridges numerous diverse DNA repair pathways^19–21^. Of particular note, mutations in nearly all FA genes cause reduced fertility, and many FA proteins are known to participate directly in or interact with meiotic DSB repair pathways^22^. FANCJ was initially identified as an interactor of BRCA1^23^, but has since been shown to interact with many other proteins including: RPA^24^, TOPBP1^25^, Bloom helicase (BLM)^26,27^, and MLH1^28–30^. Given these known interactors, many of which are critical for DSB repair events during meiotic prophase I, we hypothesized that FANCJ interacts directly with MLH1 (and perhaps other regulators) in meiosis to suppress class I CO formation.

To test this hypothesis and to further elucidate the roles of FANCJ in meiosis, we used CRISPR/Cas9 gene editing technology to generate three new *Fancj* mutant mouse lines: a complete deletion, an N-terminal mutation that deletes the MLH1 interaction site and the N-terminal region of the helicase domain, and a full length fusion gene that adds a C-terminal epitope (6xHIS-HA) to the coding sequence of *Fancj*. Simultaneously, we made a custom antibody against murine FANCJ protein, specifically targeting C-terminal residues 1124-1142. These new tools have allowed us, for the first time, to visualize FANCJ along the chromosome cores during meiotic prophase I, and have made it possible to begin probing for FANCJ protein interactors in meiosis. Although our analysis of major reproductive endpoints (sperm count and testis size) largely recapitulated our prior work, we were surprised to find no evidence in either inactivating *Fancj* mutant of an effect on MLH1 or total CO number. Our analysis further demonstrated that FANCJ and MLH1 do not interact during prophase I of meiosis and suggest FANCJ may not have any direct function in CO formation along the class I pathway. We did, however, find mild defects in DSB repair in *Fancj*-KO spermatocytes, and evidence of FANCJ colocalization with BRCA1 and TOPBP1, particularly zygonema and early pachynema. Taken together, we conclude that FANCJ has a minor, but still significant, role in early DSB processing where it may function to promote SDSA during zygonema.

## Results

### Generation of *Fancj^ΔN^, Fancj^HA^*, and *Fancj^-^* mutant mice

During meiotic studies of *Fancj^GT/GT^* males, we observed a loss of our previously reported increased meiotic recombination (as measured by the number of MLH1 foci in pachytene spermatocytes, detailed below). With the aim of both recapitulating the original phenotype and further investigating the hypothesized interaction of MLH1 and FANCJ during meiosis, we generated two novel *Fancj* inactivating mutant mice using CRISPR/Cas9. The first mutant (named *Fancj-ΔN*) featured a deletion of ∼12kb in the *Fancj* gene beginning 33bp downstream of the start codon. This deletion was predicted to remove the MLH1 interaction site (identified by Peng and colleagues^28^) and the N-terminal region of the ATP/Helicase core domain (Fig S1A). The second mutant featured a full deletion of the *Fancj* gene ranging from 277 bp upstream of the start codon through 159 bp downstream of the stop codon. Both mutants were verified by PCR genotyping of genomic ear DNA (Fig S1B) and subsequent Sanger sequencing. Likewise, PCR of testis cDNA showed expression of the *ΔN* allele, but an absence of *Fancj* transcription in *Fancj^-/-^*mice (Fig S1C).

A lack of commercially available antibodies against murine FANCJ has been a significant limitation in prior mouse studies^18,31^. Thus, we used CRISPR/Cas9 to insert a 6xHis-HA epitope tag immediately prior to the stop codon of the endogenous *Fancj* locus (which we hereafter termed *Fancj-HA;* (Fig S1A). As with the above mutants, insertion of the tag was validated by PCR genotyping of genomic ear DNA (Fig S1B). The epitope tag was further validated via Western blot, showing a band running at ∼150 kDa in testis lysate from *Fancj^HA/HA^* and *Fancj^HA/+^* mice, but no band in *Fancj^+/+^* or other mutant genotype lysate samples (Fig S1D). We additionally worked with Thermo-Fisher to produce a custom antibody against the C-terminal of the murine FANCJ (antigen sequence: DDSECFTPELFDPVDTNEE corresponding to residues 1124-1142). This antibody was validated using Western blot where, like the HA-tag, we detected a band running at ∼150 kDa in *Fancj^+/+^, Fancj^HA/HA^*, *Fancj^HA/+^*, *Fancj^ΔN/+^* and *Fancj^+/-^* testis lysates, but not in *Fancj^GT/GT^, Fancj^ΔN/ΔN^,* or *Fancj^-/-^* mutant males (Fig S1D).

### Mice homozygous for *Fancj* mutations showed reduced hematopoiesis, mild reductions in testis mass and sperm production

To assess whether our newly generated *Fancj* mutant mice display somatic phenotype consistent with prior Fanconi anemia mouse models, we started by analyzing bone marrow hematopoietic stem and progenitor cells (HSPCs) (Fig 1). *Fancj^-/-^*mice exhibit a significant loss in the number of Lineage– [Lin–] c-Kit+ Sca-1+ (LKS) HSPCs (4.7% and 1.3% for *Fancj^+/+^*and *Fancj^-/-^*, respectively, p = 0.004), harboring decreased long-term hematopoietic stem cells (LT-HSCs; 1.8% and 0.9% for *Fancj^+/+^*and *Fancj^-/-^*, respectively, p = 0.03), short-term hematopoietic stem cells (ST-HSC; 1.3% and 0.3% for *Fancj^+/+^*and *Fancj^-/-^*, respectively, p = 0.009), and multipotent progenitors (MPP; 1.3% and 0.1% for *Fancj^+/+^*and *Fancj^-/-^*, respectively, p = 0.004). The Lineage– [Lin–] c-Kit+ (LK) population was also reduced in *Fancj^-/-^* mice (25.5% and 12.9% for *Fancj^+/+^*and *Fancj^-/-^*, respectively, p = 0.004) with corresponding decreases in common myeloid progenitors (CMP; 4.2% and 1.2% for *Fancj^+/+^*and *Fancj^-/-^*, respectively, p = 0009), megakaryocytic-erythroid progenitors (MEP; 13.1% and 7.3% for *Fancj^+/+^*and *Fancj^-/-^*, respectively, p = 0.002), and granulocyte-monocyte progenitors (GMP; 5.1% and 2.3% for *Fancj^+/+^*and *Fancj^-/-^*, respectively, p = 0.017). The depletion of HSPCs in our *Fancj^-/-^* mice is in agreement with previously reported Fanconi Anemia deficient mouse phenotypes^32–35^.

**Figure 1.**
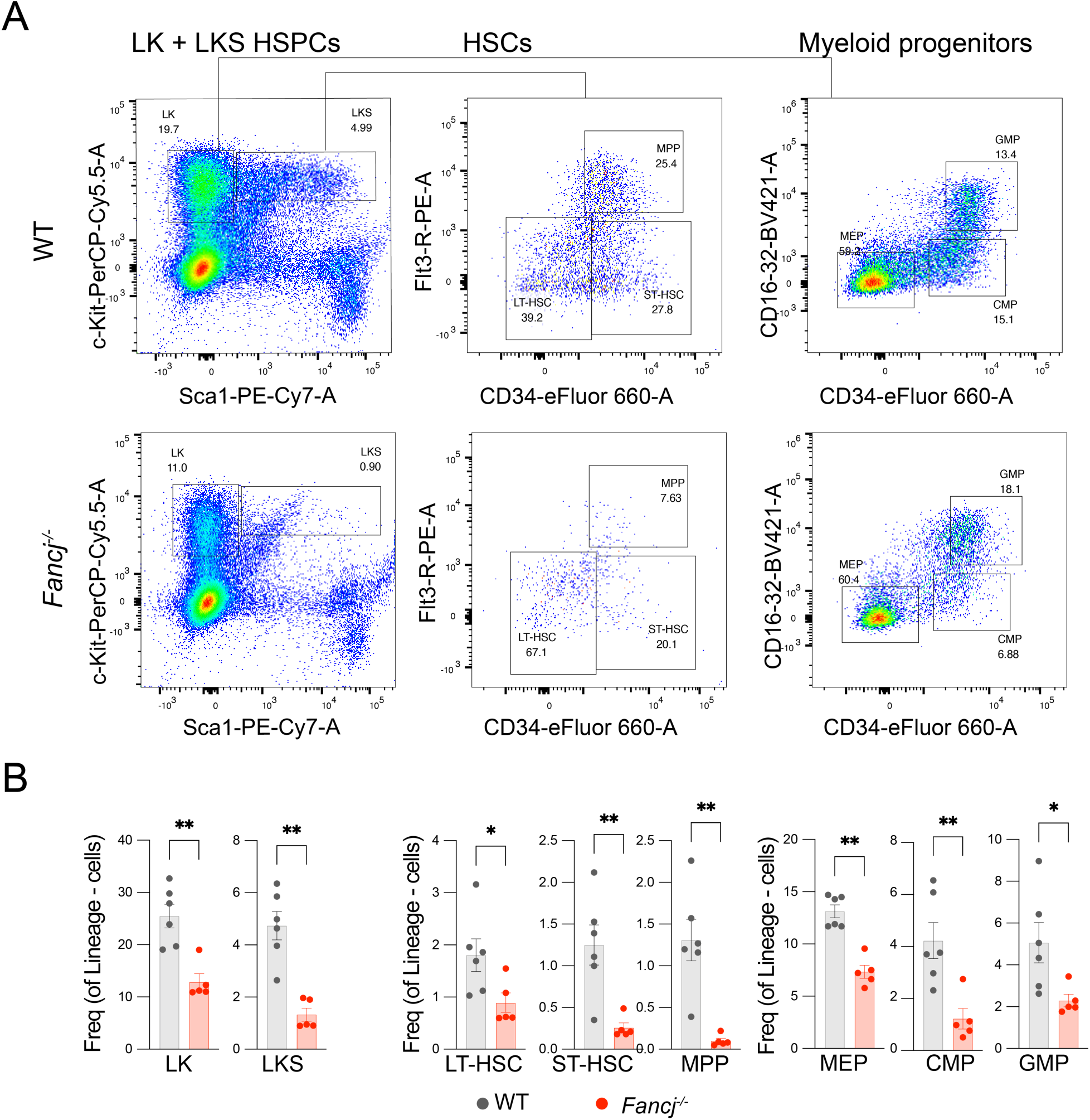
Hematopoietic stem and myeloid progenitor cells are depleted in *Fancj^-/-^* mice. Representative flow cytometry plots from *Fancj^+/+^* (WT) and *Fancj^-/-^*adult mice showing bone marrow LK, LKS, LT-HSC, ST-HSC, MPP, MEP, CMP and GMP populations. B) Quantification of the respective HSPC populations analyzed by flow cytometry in age matched *Fancj^+/+^* (black) and *Fancj^-/-^* (red) mice. Asterisks denote significant differences (* p < 0.05, ** p < 0.01).

Similar to our previous report on *Fancj^GT/GT^* mice, all *Fancj* homozygous mutant mice (*Fancj^ΔN/ΔN^*, *Fancj^-/-^*, and *Fancj^GT/GT^*) could produce viable litters. Null-by-null crosses (i.e., mating *Fancj^-/-^* males and females) did not appear to exhibit any impact on fertility and produced similar litter sizes by comparison with *Fancj^+/+^* (not shown). To assess effects of *Fancj* mutation on testis morphology, we used one-way ANOVA to compare mean testis mass as a proportion of body mass (i.e., for each mouse, mean testis mass divided by body mass) from each mutant genotype against *Fancj^+/+^* (Fig 2A; tested difference of means without assumed equal variance, Brown-Forsythe F= 11.95, p = 0.0003). Significant differences between *Fancj^+/+^* and mutant genotypes were determined by Dunnett’s multiple comparison test. Testes from *Fancj^ΔN/ΔN^*and *Fancj^-/-^* males were smaller by comparison with *Fancj^+/+^*littermate males, showing significant decreases of ∼32% and ∼34%, respectively (Fig 2A; mean ± SEM of 0.00290 ± 0.0001 [N = 13], 0.00196 ± 0.0001 [N = 9], and 0.00192 ± 0.0001 [N = 10] for *Fancj^+/+^*, *Fancj^ΔΝ/ΔΝ^* [p = 0.0012], and *Fancj^-/-^* [p = 0.0001], respectively). The magnitude of these reductions in testis mass are similar to those reported previously in the *Fancj^GT/GT^* mutant^18,31^. Although the *Fancj^GT/GT^* males in our current report also showed a slight reduction in testis mass, it was not as large as in these prior findings and, unlike the *Fancj^ΔN/ΔN^*and *Fancj^-/-^*, it did not reach the level of significance (-17% by comparison to *Fancj^+/+^*; 0.00239 ± 0.0001 [N = 8], p = 0.0684). Curiously, The *Fancj^GT/+^* showed a slight increase in testis size by comparison to *Fancj^+/+^* (∼39%; 0.004021 ± 0.0001 [N = 5], p < 0.0001). Heterozygous males for the *Fancj^ΔN^* and *Fancj^-^*alleles did not significantly differ from the *Fancj^+/+^* (mean ± SEM of 0.00309 ± 0.0001 [N = 4] and 0.00317 ± 0.0001 [N = 4] for *Fancj^ΔN/+^*and *Fancj^+/-^*, respectively).

**Figure 2.**
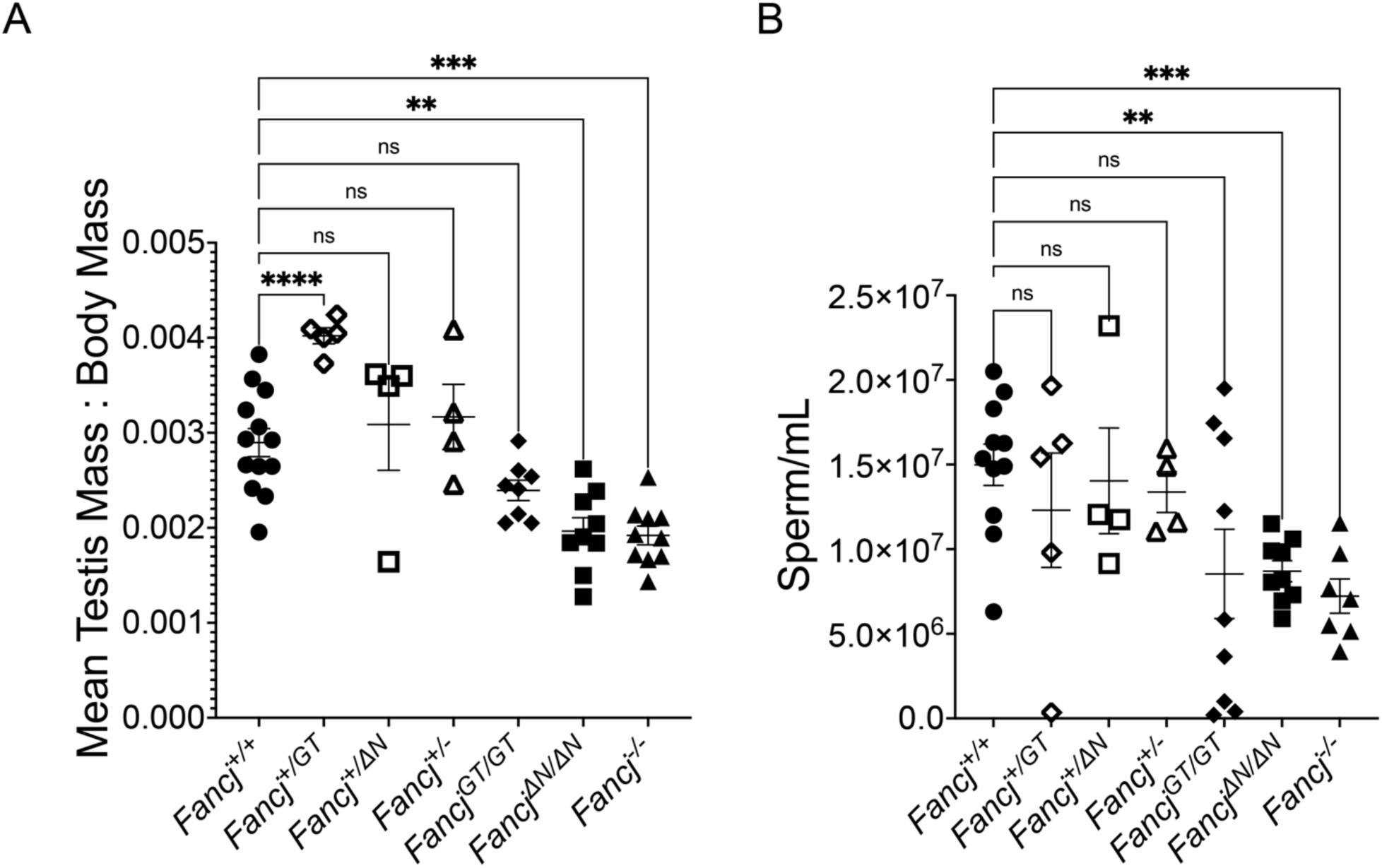
Testis mass and sperm count are significantly reduced in *Fancj* mutant males by comparison to wildtype. A) Comparison of mean testis mass as a proportion of body mass for 13 *Fancj^+/+^*, 5 *Fancj^+/GT^*, 4 *Fancj^+/ΔN^*, 4 *Fancj^+/-^*, 8 *Fancj^GT/GT^*, 9 *Fancj^ΔN/ΔN^*, and 10 *Fancj^-/-^* males. Modest reductions in testis mass were observed only in homozygous mutants, but only the *Fancj^ΔN/ΔN^* and *Fancj^-/-^* males showed a significant reduction (p = 0.0012 and p = 0.0001, respectively). B) Comparison of caudal epidydimal sperm count for 11 *Fancj^+/+^*, 5 *Fancj^+/GT^*, 4 *Fancj^+/ΔN^*, 4 *Fancj^+/-^*, 9 *Fancj^GT/GT^*, 9 *Fancj^ΔN/ΔN^*^,^ and 7 *Fancj^-/-^* males. As with testis mass, sperm counts were reduced only in homozygous mutants, with only the *Fancj^ΔN/ΔN^* and *Fancj^-/-^* showing significant reduction (p = 0.0021 and p = 0.001, respectively). Asterisks denote significant differences (* p< 0.05, ** p < 0.01, *** p < 0.001, **** p < 0.0001).

To assess whether *Fancj^ΔΝ/ΔΝ^* and *Fancj^-/-^* male mice show similar reductions in epididymal sperm as that reported for *Fancj^GT/GT^* males, we used a one-way ANOVA to compare caudal total epididymal spermatozoa counts from each mutant genotype against *Fancj^+/+^* (Fig 2B; Brown-Forsythe F=1.687, p=0.0465). Significant differences between the *Fancj^+/+^*mean sperm count and that of each mutant genotype were determined by Dunnett’s multiple comparison test. Heterozygous males for the *Fancj^ΔN^* and *Fancj^-^* alleles showed no difference in sperm count to that of their *Fancj^+/+^* littermate control males (Fig 2B). Similar to our observations of testis mass, both *Fancj^ΔΝ/ΔΝ^* and *Fancj^-/-^* males showed significant reductions in sperm production of ∼42% and ∼52%, respectively, by comparison with the *Fancj^+/+^* (Fig 2B, p=0.0021 and 0.001, respectively). Although the average sperm count of *Fancj^GT/GT^* males was nearly equivalent to that of *Fancj^ΔΝ/ΔΝ^* males, there was significantly higher variation among the *Fancj^GT/GT^* males, precluding them from reaching statistical significance.

### FANCJ localizes as discrete foci on chromosome cores beginning in leptonema of prophase I

To characterize the distribution of FANCJ throughout meiotic prophase I, we performed indirect immunofluorescence staining of spermatocyte chromosome spreads using our custom antibodies against FANCJ and the axial element (AE) of the synaptonemal complex, SYCP3 (Fig 3A). FANCJ first appears as discrete foci diffusely localized throughout the nucleus of leptotene spermatocytes, with many dim foci localizing to nascent SYCP3 filaments. The most prominent localization pattern for FANCJ occurred during mid-zygonema when hundreds of bright foci localized along the lengths of SYCP3 AEs. By late zygonema, discrete foci of FANCJ could scarcely be distinguished, rather appearing as a filament co-localizing with most unsynapsed SYCP3. This localization pattern persisted into pachynema where FANCJ localized exclusively to the unsynapsed SYCP3 of the X and Y chromosomes (Fig 3Ei). The XY localization of FANCJ could be seen in diplonema, but no clear FANCJ could be observed in diakinesis (not shown). To validate these results, we also stained *Fancj^-/-^* spermatocyte chromosome spreads with antibodies against FANCJ and SYCP3 (Fig 3C). The *Fancj^-/-^*spermatocyte nuclei lacked any FANCJ signal from leptonema through zygonema; however, some staining was visible on the unsynapsed regions of the X and Y chromosomes in pachytene, similar to that observed in *Fancj^+/+^*(Fig 3Eiii). Although this signal was gone in the *Fancj^-/-^*by diplonema, its presence in the *Fancj^-/-^* raised the possibility that the XY localization in pachynema was a non-specific target of our antibody.

**Figure 3.**
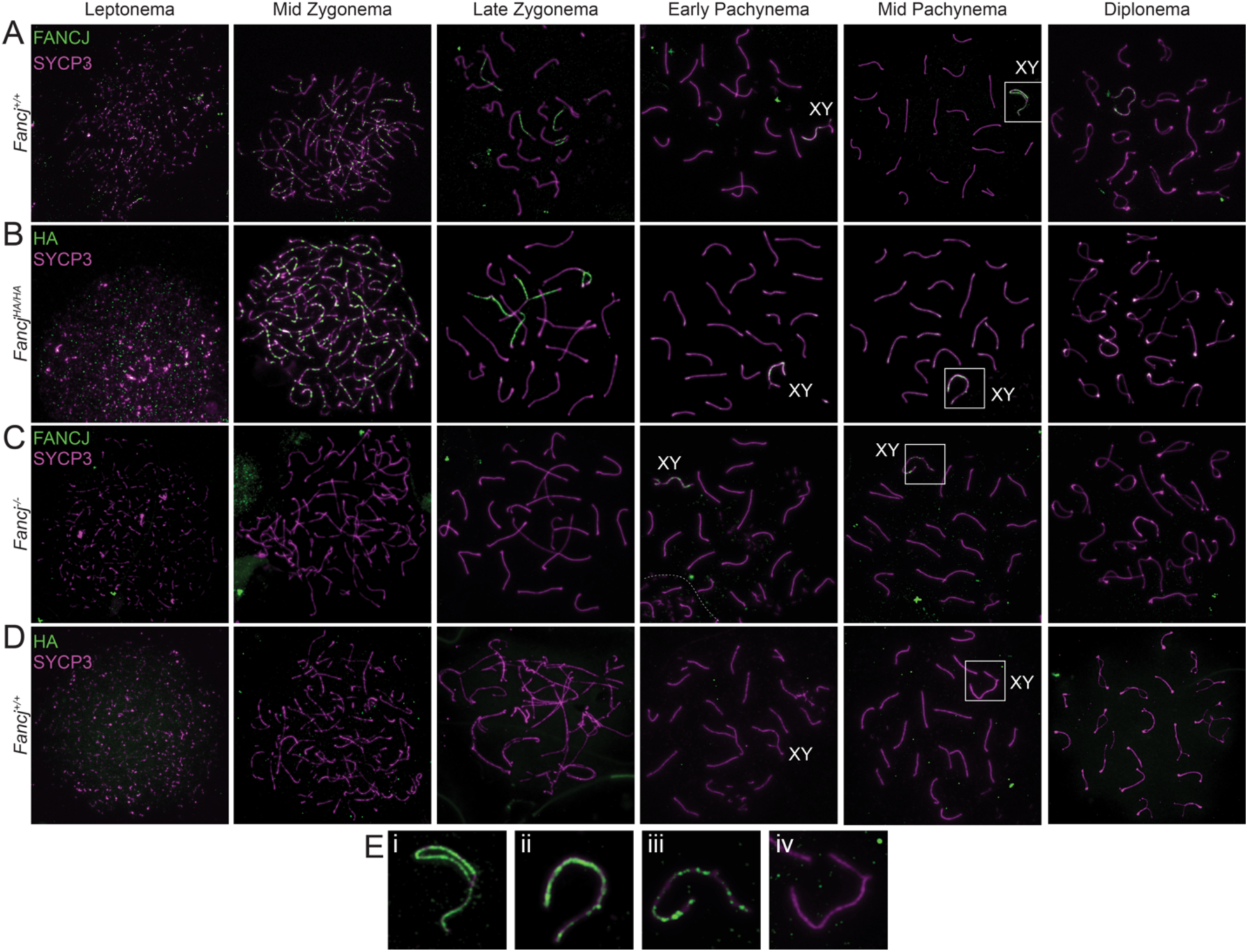
FANCJ localization on chromosome axes during meiotic prophase I. Immunofluorescence imaging across meiotic prophase I are shown. FANCJ (green) and SYCP3 (magenta) were co-stained in *Fancj^+/+^* (A) *Fancj^-/-^* (C), with the latter as a negative control. Similarly, HA (green) and SYCP3 (magenta) were co-stained in *Fancj^HA/HA^* (B) and *Fancj^+/+^*(D), with the latter as a negative control. Both FANCJ and the HA-tagged endogenous FANCJ protein localized on chromosome axes (SYCP3) as discrete foci beginning in leptonema-mid zygonema, and then as dense accumulations along unsynapsed SYCP3 axes in late zygonema. All leptotene and zygotene signal for the FANCJ and HA antibodies appear to be specific, as neither was found in their respective negative controls. Although the FANCJ antibody also appears on the XY chromosomes in early pachynema and persists into diplonema, some of this signal appears to be non-specific and the FANCJ antibody also appears on the XY of the *Fancj^-/-^*negative control. The HA antibody also appeared on the XY but was less saturated than the FANCJ antibody and did not persist as long. Further, the HA antibody did not show any signal on the XY of the negative control, suggesting FANCJ localizes to the XY in early pachynema and begins to dissociate in mid-pachynema. Zoomed-in images of the XY in mid-pachynema are shown in E) panels for FANCJ in *Fancj^+/+^* (i), HA in *Fancj^HA/HA^* (ii), FANCJ in *Fancj^-/-^* (iii), and HA in *Fancj^+/+^* (iv).

To clarify the observations made using our custom FANCJ antibody, we also examined FANCJ localization throughout prophase I via indirect immunofluorescence of *Fancj^HA/HA^* spermatocytes using antibodies against the HA epitope tag (Fig 3B). Staining against HA largely replicated that of the FANCJ antibody; however, HA localization on the XY (Fig 3Eii) did not appear to persist into diplonema. Instead, signal on the XY appears to diminish after early pachynema and could rarely be seen in late pachytene spermatocytes (not shown). Staining *Fancj^+/+^*spermatocytes with antibodies against the HA epitope tag showed the expected absence of signal in each stage of prophase I (Fig 3D, Eiv).

To further confirm that antibodies against the FANCJ-HA epitope tag and our custom FANCJ antibody detect the same protein, we co-immunostained *Fancj^HA/HA^* spermatocytes with antibodies to HA and FANCJ. Dual immunofluorescence with both antibodies as primary antibodies, using different secondary antibodies (AF488 and AF594, respectively) showed strong co-localization in zygonema and pachynema (Fig 4A-E). These data collectively support a FANCJ localization pattern that begins in leptonema, culminates as bright, SYCP3-associated discrete foci in zygonema, is restricted to unsynapsed SYCP3 regions in late zygonema, and finally is exclusive to the XY until mid/late pachynema.

**Figure 4.**
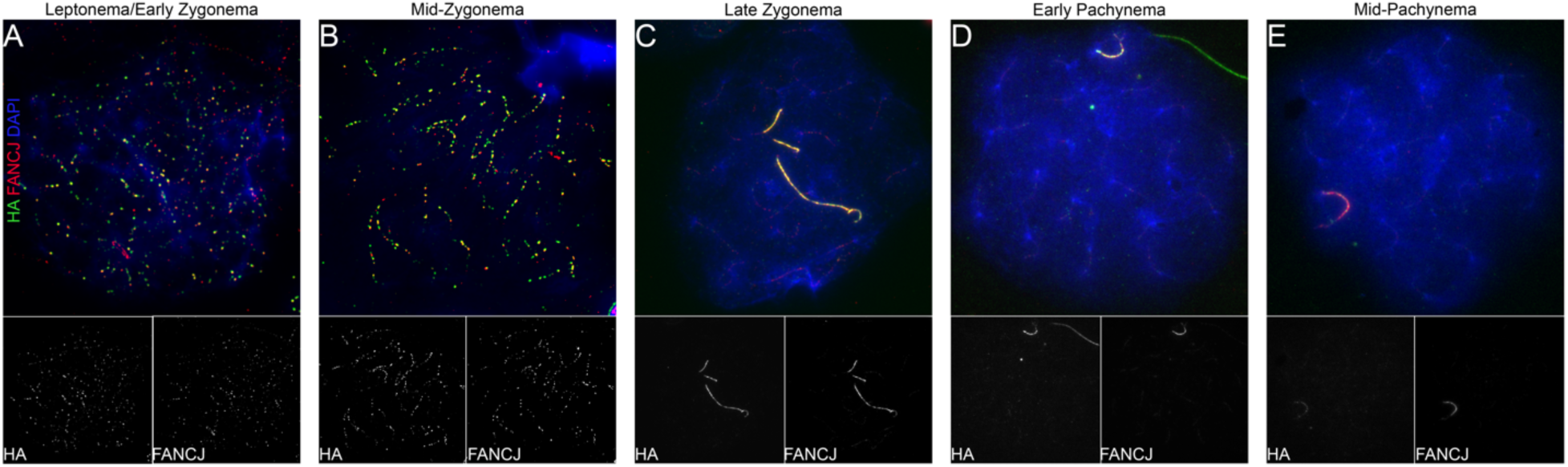
FANCJ and FANCJ-HA co-localize. *Fancj^HA/HA^* spermatocytes in leptonema/early zygonema (A), mos-zygonema (B), late zygonema (C), early pachynema (D), and mid-pachynema (E) immunostained with antibodies to HA (green), FANCJ (red), and DAPI (blue). Individual HA and FANCJ channels are shown in greyscale below their respective composite image.

### Loss of *Fancj* impairs DSB repair in meiotic prophase I

To determine the effects of FANCJ loss on meiotic prophase I progression, we evaluated whether spermatocytes from *Fancj^-/-^* males showed defects in DSB repair. We used immunofluorescence staining of the RecA homolog, RAD51, as a proxy for DSB induction and initial processing. RAD51 localizes to DSBs and facilitates DMC1 filament assembly, thus promoting strand invasion during meiotic recombination^6–8^. During typical prophase I in mouse spermatocytes, hundreds of RAD51 foci accumulate on the chromosome cores in zygonema, but by pachynema RAD51 is localized almost exclusively to the unsynapsed regions to the XY, while loss of RAD51 on autosomes signifies DSB repair progression beyond the strand invasion steps. We thus quantified RAD51 total foci number in zygonema (Fig 5A-C) and autosomal foci in pachytene spermatocytes (Fig 5D-F) from *Fancj^+/+^* and *Fancj^-/-^* mice and determined significant differences using a two-tailed t-test. There was no difference in RAD51 focus counts between *Fancj^-/-^*(mean ± SEM of 214.9 ± 4.5) and *Fancj^+/+^*(mean ± SEM of 206.6 ± 5.5) spermatocytes in zygonema (Fig 5C; t = 1.171, p = 0.245). However, *Fancj^-/-^* males had significantly more autosomal RAD51 foci in pachynema by comparison to that observed in *Fancj^+/+^* males (Fig 5F; mean ± SEM of 23.9 ± 2.0 and 5.9 ± 0.6, respectively; t = 8.764, p < 0.0001) indicative of delayed or defective DSB repair.

**Figure 5.**
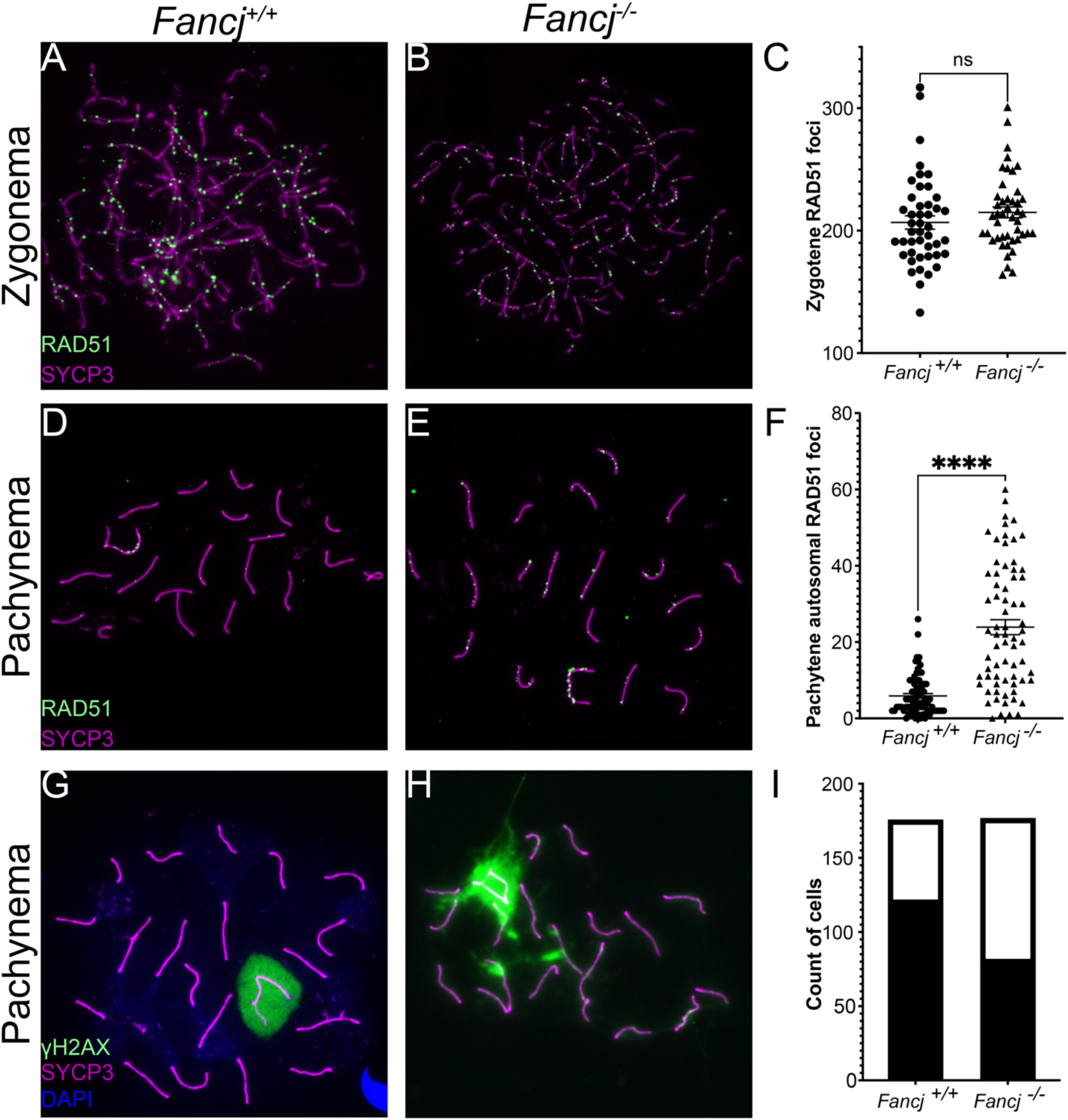
*Fancj*-/- males showed a significant increase in persistent DSBs in pachynema. *Fancj^+/+^* zygotene (A) and pachytene (D) and *Fancj^-/-^* zygotene (B) and pachytene (E) spermatocytes were immunostained with antibodies to RAD51 (green) and SYCP3 (magenta). C)There was no significant difference in the number of RAD51 foci between *Fancj^+/+^* (2 males, 45 cells total) and *Fancj^-/-^* (2 males, 45 cells total) in zygotene). F) There was significantly more autosomal RAD51 in *Fancj^-/-^* pachytene spermatocytes (3 animals, 70 cells total) by comparison with *Fancj^+/+^*(3 animals, 73 cells total). Pachytene spermatocytes from *Fancj^+/+^*(G) and *Fancj^-/-^* (H) mice were immunostained with antibodies to γH2AX (green) and SYCP3 (magenta). *Fancj^-/-^* mice (4 animals,176 cells total) showed significantly more aberrant γH2AX localization (i.e., autosomal localization) by comparison to the *Fancj^+/+^* (4 animals, 175 cells total). I) Bar graph shows number of cells with normal (black) and atypical (white) γΗ2ΑΧ localization.

To confirm a DSB repair defect in *Fancj^-/-^* males, we performed immunofluorescence staining of the phosphorylated Ser139 form of H2A histone family member X (γH2AX), a well-established marker of DSBs^36^. In typical spermatocyte meiosis, γH2AX signal is diffuse throughout the nucleus in leptonema and zygonema, where it marks sites of DSB induction, but is restricted specifically to the XY-bearing sex body in pachynema and diplonema as a part of meiotic chromosome sex inactivation (MSCI)^37–39^. Any non-sex body associated γH2AX signal in pachynema and diplonema reflects delayed or aberrant DSB repair^40–42^. Cells with any γH2AX signal outside the sex body, or with flares emanating from the sex body, during pachynema were classified as “atypical”, while cells displaying a γH2AX signal localized only in the sex body were classified as “normal”. The total number of cells in each category for *Fancj^+/+^*and *Fancj^-/-^* males were compared by Fisher’s exact test. *Fancj^-/-^* males had significantly fewer cells with normal γH2AX signal by comparison to *Fancj^+/+^* (Fig 5G-I; 46% and 69%, respectively, p < 0.0001). Collectively, the atypical γH2AX signal and persistent RAD51 foci at pachynema in *Fancj^-/-^* spermatocytes suggest that FANCJ is necessary for correct and timely progression of DSB repair in leptonema and zygonema.

### Crossover formation is unaffected in *Fancj* mutant mice

In somatic cells, FANCJ is reported to interact with the mismatch repair protein MutL homolog 1 (MLH1) to ensure correct interstrand crosslink (ICL) repair^28–30,43^. In meiosis, MLH1 is the final designation marker of virtually all COs, i.e., class I COs (∼90-95% of all COs)^12,13^. That FANCJ and MLH1 interact in somatic cells led us to hypothesize that FANCJ plays a role in regulating CO formation through such an interaction. To test this hypothesis, we quantified the number of class I COs (measured as MLH1 foci in pachytene spermatocytes; Fig 6A, B) and compared the mean number from each *Fancj* mutant (*Fancj^ΔN/ΔN^*, *Fancj^-/-^*, and *Fancj^GT/GT^*) genotype to *Fancj^+/+^*by one-way ANOVA (F = 2.792, p = 0.01). Significant differences between *Fancj^+/+^* and mutant genotypes were determined by Dunnett’s multiple comparison test. Notably, neither *Fancj^ΔΝ/ΔΝ^*nor *Fancj^-/-^* males showed a significant difference in MLH1 foci number at pachynema either (Fig 6C; mean ± SEM of 24.1 ± 0.2, p = 0.999 and 23.9 ± 0.3, p = 0.996, respectively). When we examined *Fancj^GT/GT^* to confirm that we saw the same effect, we found the MLH1 counts from *Fancj^GT/GT^* males also did not show any difference to that of *Fancj^+/+^* littermate control males (Fig 6C; mean ± SEM of 23.5 ± 0.2 and 24.1 ± 0.2, respectively, p = 0.187). This is in contrast to our prior report that the gene trap disruption of *Fancj* caused a significant increase in MLH1 foci^18^. Collectively, these data suggest none of the *Fancj* mutants affect class I CO number in mouse spermatocytes.

**Figure 6.**
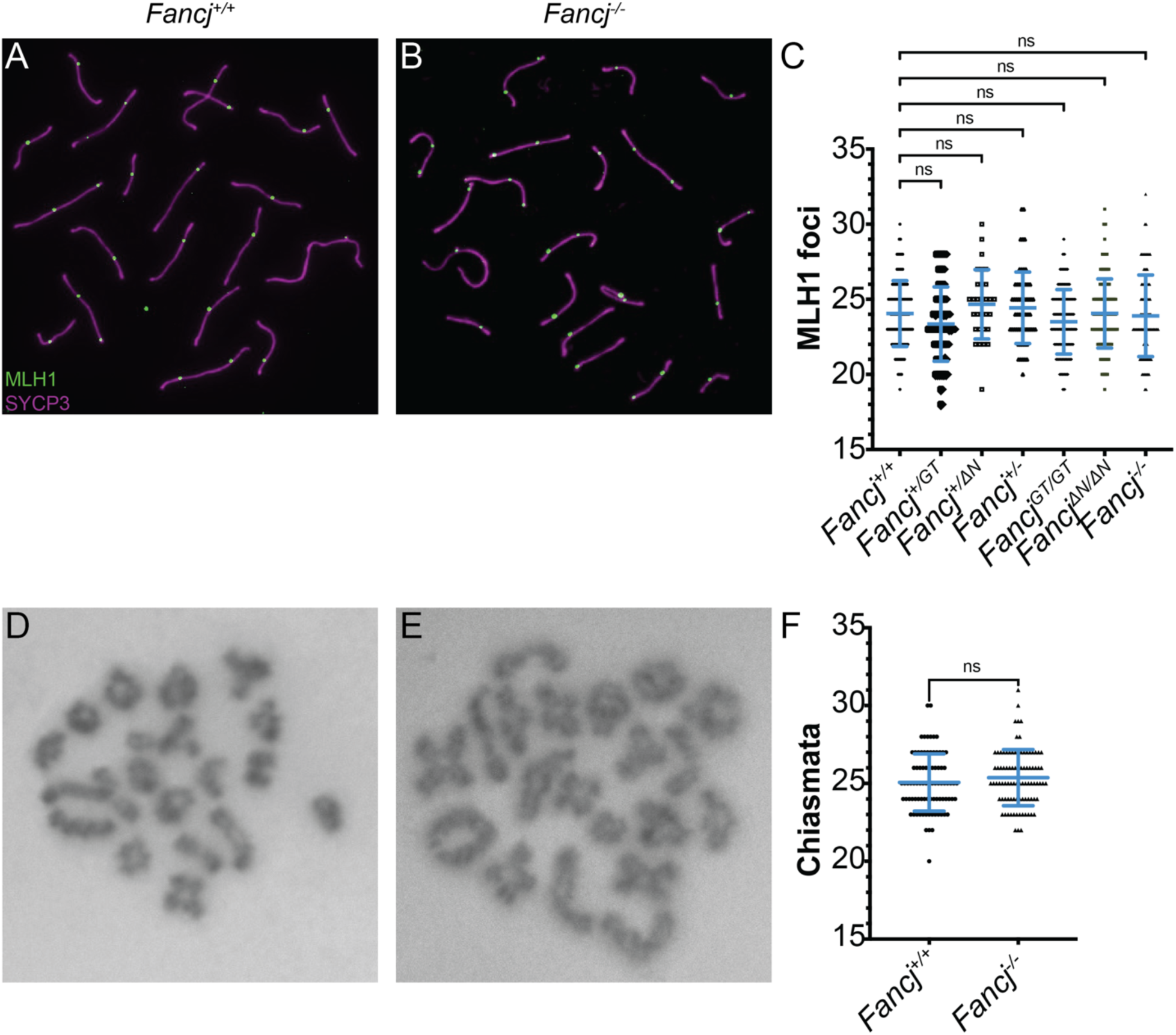
*Fancj* mutants show no significant change in MLH1 foci or total crossover number. Pachytene spermatocytes from *Fancj^+/+^* (A) and *Fancj^-/-^* (B) mice immunostained with antibodies to MLH1 (green) and SYCP3 (magenta). C) Comparison of mean MLH1 foci number for 4 *Fancj^+/+^*, 2 *Fancj^+/GT^*, 1 *Fancj^+/ΔN^*, 2 *Fancj^+/-^*, 7 *Fancj^GT/GT^*, 5 *Fancj^ΔN/ΔN^*^,^ and 3 *Fancj^-/-^* males (25-30 cells analyzed per male).Chiasmata stained with Giemsa in diakinesis from *Fancj^+/+^* (D) and *Fancj^-/-^* (E) mice. F) Comparison of mean chiasmata counts for *Fancj^+/+^* and *Fancj^-/-^* mice (3 males per genotype, 25-30 cells analyzed per male).

To determine whether loss of *Fancj* had any impact on total CO number, we also quantified the number of chiasmata (the physical manifestation of COs) in spermatocyte diakinesis of *Fancj^+/+^* and *Fancj^-/-^* mice (Fig 6D, E). Significant difference from the *Fancj^+/+^* mean was determined using a two-tailed t-test. *Fancj^+/+^* males had 25.1 ± 0.2 chiasmata per cell (mean ± SEM) while *Fancj^-/-^*males had 25.4 ± 0.2 chiasmata per cell showing no significant difference between the two genotypes (Fig 6F; t = 1.117, p = 0.84).

### FANCJ does not colocalize or interact with key pro-crossover factors in later stages of prophase I

To assess whether FANCJ associates with proteins that stabilize recombination intermediates to promote CO formation (i.e., pro-CO factors) in meiosis, we performed co-immunofluorescence staining *Fancj^HA/HA^* spermatocyte chromosome spreads with antibodies against the HA epitope tag, MSH4, MLH1, and MLH3 (Fig 7A-E). MSH4 (along with MSH5) is a component of the mismatch repair heterodimer, MutSγ. It is thought MutSγ binds to CO intermediates (e.g., Holliday junctions), and that MutLγ (MLH1 and MLH3) is recruited to a subset of MutSγ sites to produce class I COs^12,14,15^. We previously found that disruption of MutSγ function eliminates all chiasmata, suggesting normal MutSγ function is important for all CO formation and not just those of the class I variety.^44^ We observed some colocalization of FANCJ-HA with MSH4 in mid-zygonema, decreasing in late zygonema (Fig 7A); however, by pachynema, the respective signals of FANCJ-HA and MSH4 appeared to be almost completely mutually exclusive (Fig 7B-D). FANCJ-HA localized neither to sites of maturing class I COs in mid-pachynema (i.e., marked by both MSH4 and MLH3; Fig 7D) nor to the ultimate sites of class I COs marked by MLH1 and MLH3 (Fig 7E). Collectively, these data suggest FANCJ does not localize to CO intermediates in prophase I.

**Figure 7.**
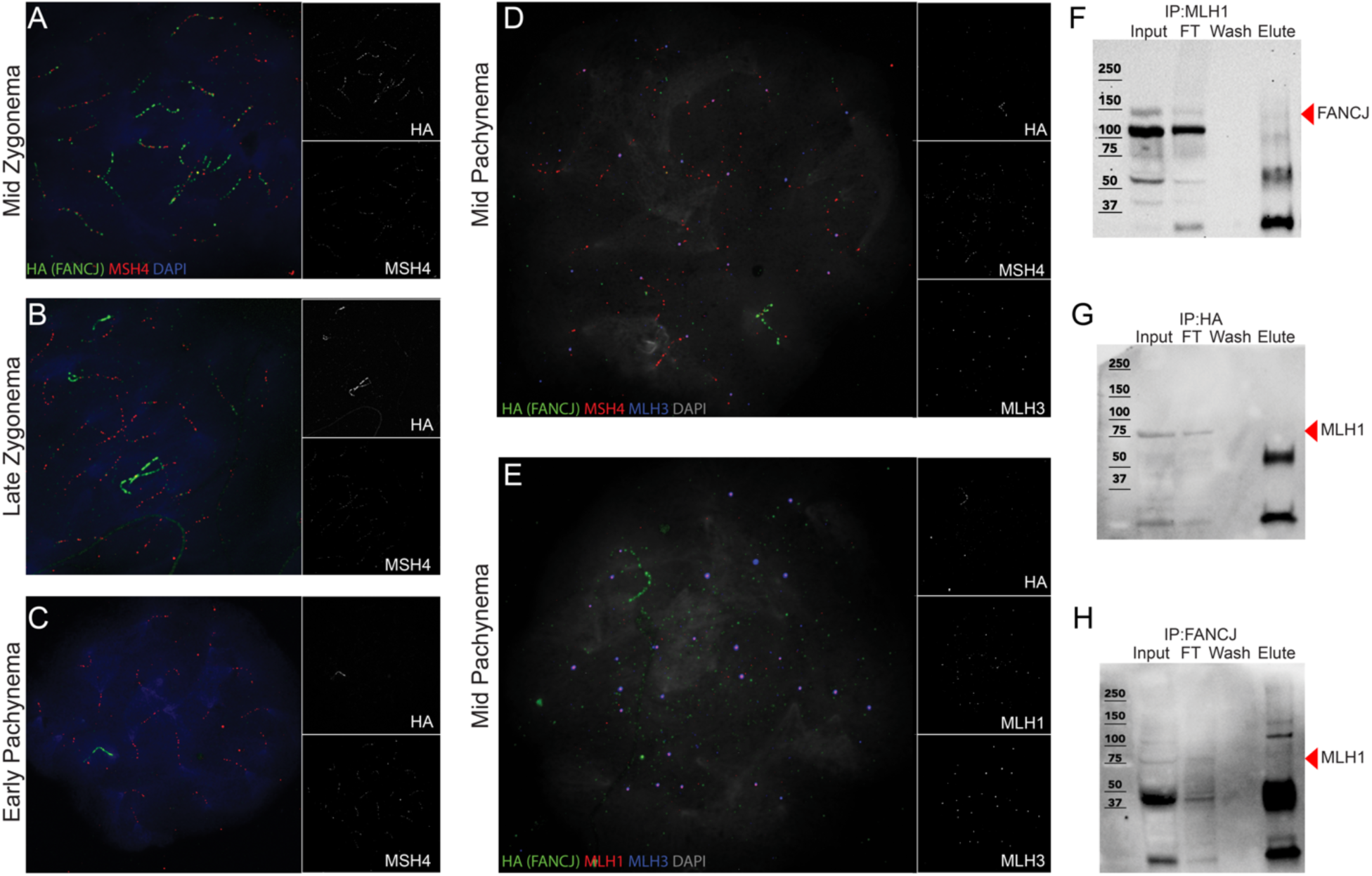
FANCJ does not co-localize with MSH4/MLH3 or interact with MLH1 in late prophase I. Spermatocytes from *Fancj^HA/HA^* mice showing an absence of FANCJ colocalization with MSH4, MLH3, and MLH1. A) Mid-zygotene, B) late zygotene, and C) early pachytene cells immunostained with antibodies to HA (FANCJ; green), MSH4 (red), and DAPI (blue). Mid-pachytene cells immunostained with antibodies to D) HA (green), MSH4 (red), MLH3 (blue), and DAPI (grey) and E) HA (green), MLH1 (red), MLH3 (blue), and DAPI (grey). Individual greyscale channels for HA, MSH4, MLH3, and MLH1 are besides their respective composite images. F) Western blot of MLH1-IP using whole testis lysate input. Sample was run on an 8% SDS-PAGE gel, and membrane blotted with antibody against HA. Black arrow highlighting FANCJ bands (expected size 131 kDa but detected at ∼150 kDa). G) Western blot of HA-IP using *Fancj^HA/HA^* whole testis lysate input. Sample was run on an 8% SDS-PAGE gel, and membrane blotted with antibody against MLH1. Black arrow highlighting MLH1 bands (84.6 kDa). H) Western blot of HA-IP using whole testis lysate input. Sample was run on an 8% bis-acrylamide gel, and membrane blotted with antibody against MLH1. Black arrow highlighting MLH1 bands (84.6 kDa).

To confirm that FANCJ does not associate with MLH1, as suggested by the temporally distinct localization of each protein (Fig 7E), we performed anti-MLH1 immunoprecipitation (IP) followed by western blot against FANCJ using *Fancj^+/+^* whole testis lysate (Fig 7F). We found that MLH1 did not pull down FANCJ, although FANCJ was detected in both the input and the flow-through (FT). Similarly, MLH1 was not detected in the elution of reciprocal anti-HA-IP (Fig 7G) and anti-FANCJ-IP (Fig 7H), but MLH1 was present in the input and FT of each. MLH1 was detected in the anti-MLH1-IP elution (Fig S2A); similarly, FANCJ was detected in both the anti-HA-IP (Fig S2B) and anti-FANCJ-IP (Fig S2C) elutions, suggesting the immunoprecipitation experiments were successful. As further validation that we could detect an MLH1-FANCJ interaction in other DNA repair contexts^28,30^, we performed an anti-FANCJ-IP using irradiated (IR) and untreated (No-IR) mouse pre-B cell lysates (Fig S3). In both instances, MLH1 was observed in the anti-FANCJ-IP elution and not the anti-IgG-IP negative control. Thus, although we can confirm that FANCJ interacts with MLH1 in the context of some somatic DNA repair events, there appears to be no such interaction between FANCJ and MLH1 in mouse spermatocytes.

### Mass spectrometry of FANCJ and FANCJ^HA^ could not identify specific FANCJ interactors

To identify putative FANCJ interactors in meiosis, we initially prepared three biological replicates consisting of age-matched whole testis and thymus lysates that were used for IP against our custom rabbit anti-FANCJ antibody and a rabbit IgG negative control. IP elutions were submitted for label free mass spectrometry (MS). A total of 1171 proteins were identified (Table S1), and this number was further refined to 187 proteins that were identified by at least 10 peptides. The list of putative interactors was further narrowed using an abundance ratio (based on the peptide spectrum match (PSM) number) of FANCJ-IP:IgG-IP of at least 10, resulting in a list of 21 proteins, none of which had previously been characterized as an interactor of FANCJ. Included were proteins involved in DNA repair (e.g., ATRIP, TEX11, RUVBL1/2, and USP4), chromatin organization (e.g., TRIM28), and RNA processing (e.g., ADAD2, RNF17, and MOV10). To validate these findings, we performed anti-FANCJ-IP on whole testis lysate from *Fancj^+/+^* and *Fancj^-/-^* mice, and followed with western blot using antibodies against ATRIP, RUVBL1, USP4, TRIM28, and ADAD2 (the five most promising putative interactors based on abundance ratio and peptide number; Fig S4). TRIM28 was detected in all sample inputs, but not in any elution, ruling it out as an interactor (Fig. S4F). ATRIP, RUVBL1, USP4, and ADAD2 were detected in the *Fancj^+/+^*anti-FANCJ-IP elution but not the anti-IgG-IP negative control; however, all four were also detected in the anti-FANCJ-IP elution from *Fancj^-/-^*testes (Fig S4B-E). Subsequent anti-HA-IP followed by WB from *Fancj^HA/HA^* and *Fancj^+/+^* testis lysates confirmed that all five proteins were likely non-specific and not true interactors of FANCJ.

We next performed IP-MS using anti-HA antibodies on protein extracts from enriched germ cells of *Fancj^HA/HA^* and *Fancj^+/+^* (negative control) age-matched mice, with a total of two biological replicates. Proteins eluted from anti-HA beads were digested with trypsin and the resulting peptides were labelled with tandem mass tag (TMT) reagent. Labelled peptides recovered from *Fancj^HA/HA^* and *Fancj^+/+^* IPs were combined and fractionated by HILIC chromatography to increase coverage of detected and quantified peptides. As shown in Figure 8 and Table S2, analysis of both biological replicates revealed a significant enrichment of FANCJ in the *Fancj^HA/H^* IP as well as significant enrichment of CLASR, a putative regulator of splicing that was not previously reported to interact with FANCJ. We did not detect proteins previously reported to interact with FANCJ, including TOPBP1 or BRCA1, in the elutions. Notably, the number of FANCJ peptides detected in these analyses was low (6 in replicate 1 and 14 in replicate 2). Therefore, the inability to detect known interactors of FANCJ is likely attributed to the low abundance of the bait protein, making it technically challenging to detect interacting proteins that are expected to be present at lower abundance in the protein elutions.

**Figure 8.**
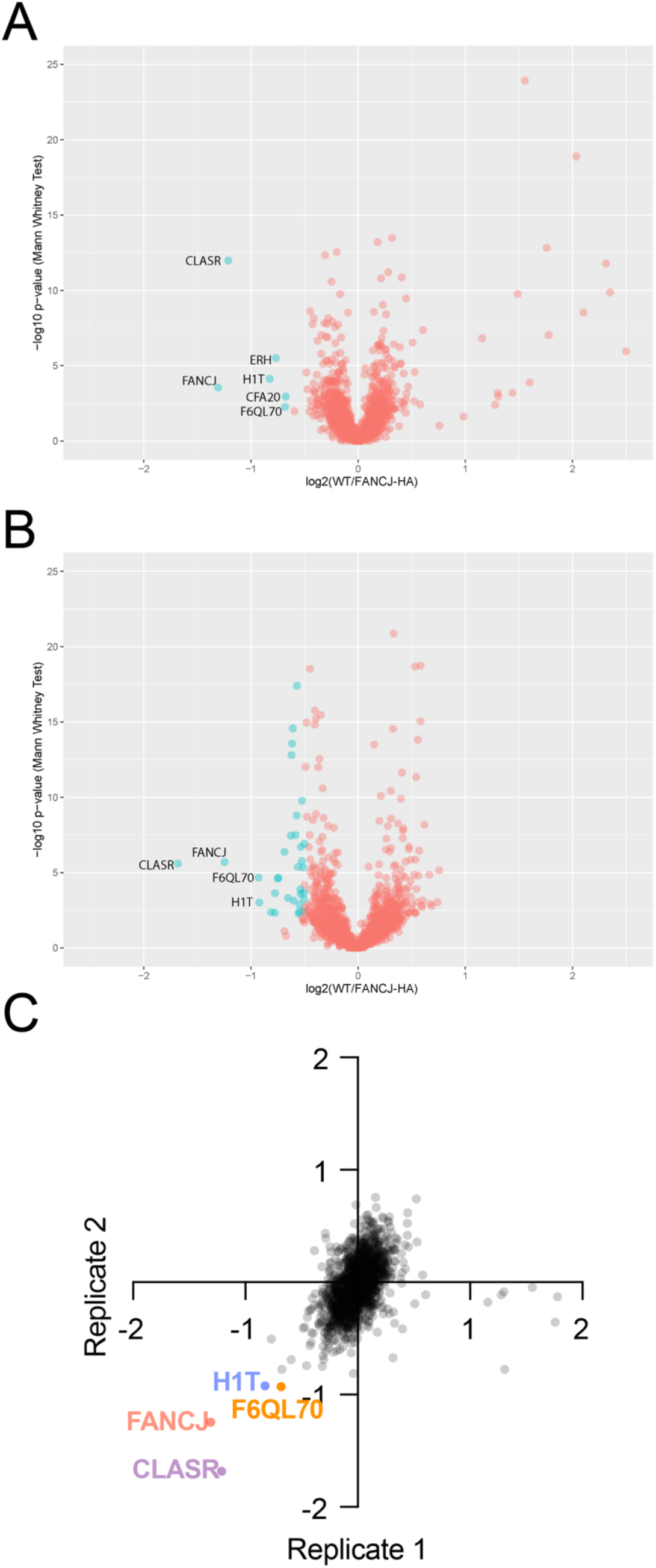
FANCJ-HA IP-MS volcano plots shows possible interactors. The first (A) and second (B) biological replicates of the αHA IP-MS performed on enriched germ cell lysates from *Fancj^HA/HA^* and *Fancj^+/+^* mice. Points of the positive x-axis correspond to proteins enriched in the *Fancj^+/+^* negative control sample; points on the negative x-axis correspond to proteins enriched in the *Fancj^HA/HA^*sample. Potential FANCJ interactors highlighted in cyan and non-interactors in red. Notably, the total abundance of FANCJ in each replicate is strikingly low, making it difficult to discern whether the identified interactors are likely to be real. C) Scatter plot of IP-MS/MS datasets corresponding to WT/FANCJ-HA, first IP (Y-axis) and WT/FANCJ-HA, second IP (X-axis) from total germ cells. Quadrant 4 displays the potential interactors that appeared on both replicates.

### FANCJ strongly colocalizes with BRCA1 and TOPBP1 throughout meiotic prophase I

Having pushed IP-MS as far as we could, we opted to use co-immunofluorescence staining of prophase I chromosome spreads to identify proteins that co-localized with FANCJ-HA. Although we were limited by antibody availability, we were able to co-stain using antibodies against BRCA1 and TOPBP1. Both BRCA1 and TOPBP1 are reported to interact with FANCJ in somatic cells^23,25^ and both are involved in the establishment of ATR signaling in response to programmed DSB induction at the onset of meiotic prophase I^45^. In early/mid zygonema, BRCA1 and TOPBP1 each partially co-localized with FANCJ-HA (Fig 9A, E). Beginning in late zygonema, both BRCA1 and TOPBP1 strongly co-localized with FANCJ-HA and their signals almost perfectly overlapped (Fig 9 B-C, F-G). This level of strong localization persisted until mid-pachynema, when TOPBP1 began to load on the chromatin loops of the sex body (Fig 9H). Although FANCJ-HA does still localize to the cores of the XY in mid-pachynema (much like TOPBP1 and BRCA1), the signal intensity of FANCJ-HA began to wane, and its localization to the XY did not persist as long as that of BRCA1 and TOPBP1 (Fig 9D, H). The antibody we used against BRCA1 did not perform well on western blots, precluding evaluation of a meiotic FANCJ-BRCA1 interaction by IP-WB. However, we were able to perform an anti-HA-IP from *Fancj^HA/HA^* and *Fancj^+/+^* whole testis lysate and did not detect TOPBP1 in the elution (Fig S5). This suggests that while TOPBP1 and FANCJ localize to the same sites and the same time during meiosis, their interaction is possibly indirect or at too low abundance to be detected by IP-WB or other biochemical means.

**Figure 9.**
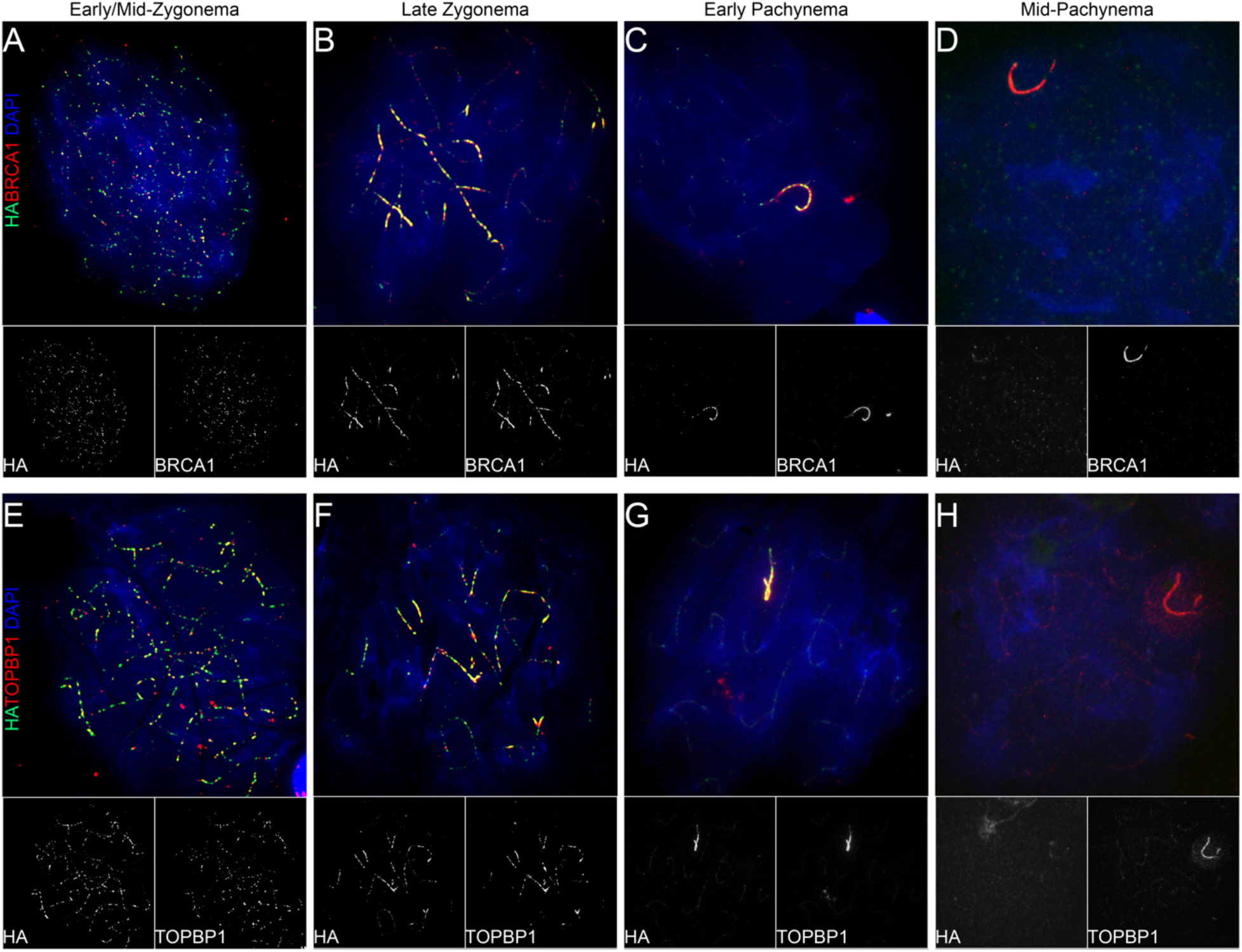
FANCJ shows strong co-localization with BRCA1 and TOPBP1. *Fancj^HA/HA^* early/mid and late zygotene and early and mid-pachytene spermatocytes were immunostained with antibodies against HA (FANCJ; green) and in red either BRCA1 (A-D) or TOPBP1 (E-H), and DAPI. Greyscale images of individual HA, BRCA1 and TOPBP1 channels are below their respective composite image.

## Discussion

The current analysis of FANCJ function in meiosis represents the most complete investigation to date of this poorly understood FA protein in mammalian meiosis. To elucidate the function of FANCJ in the mouse, we created two novel inactivating mutant mouse lines, along with a dual epitope-tagged mouse allele, and a novel custom antibody. The full deletion of *Fancj* elicited very subtle meiotic phenotypes overall, making it difficult to discern a mechanistic function. Although most FA mutations present with significant infertility or subfertility^22^, we found loss of *Fancj* to have little impact on overt viability or fertility. Consistent with previous findings^18,31^, both the *Fancj* inactivating mutant mouse lines showed significant decreases in testis size and sperm count, but all homozygous mutants were fertile. Previous observations of the gene-trap *Fancj* disruption pointed to increased prospermatogonial death, but no significant increase in germ cell loss after 7 days post-partum^18^. Although we did not examine the perinatal testis in our current study, these previous findings point to a likely roll for FANCJ in DNA repair in the rapidly expanding neonatal germ line that likely would be congruous with its function in somatic replication-coupled DNA repair. An important distinction here is our current study suggests FANCJ has similar yet distinct functions in meiotic DNA repair.

Studies of FANCJ in somatic cells have described a diversity of functions in DNA repair, including facilitating repair of ICLs via homologous recombination (HR) and resolving aberrant structures impeding DNA replication, e.g., G-quadruplex (G4) DNA^46,47^ and DNA-protein crosslinks^48^. Such studies have also uncovered several interactors of FANCJ, including BRCA1^23^, MLH1^28^, MSH5^49^, BLM^26,27^, MRE11^50^, CtIP^51^, RPA^24^, and TOPBP1^25^. This complexity of roles and possible interactors makes it difficult to elucidate FANCJ function in mammalian meiosis. We hypothesized that due primarily to its interaction with MLH1 and BLM, FANCJ likely had a role in regulating CO formation. Specifically, FANCJ might prevent MLH1 accumulation at joint molecules and work with BLM to produce NCOs. Our previous report of gene-trap disruption of *Fancj* supported this hypothesis, as mice homozygous for the gene-trap showed elevated MHL1 foci in pachynema and increase COs in diakinesis^18^. However, our current findings contradict previous reports, finding no change in MLH1 focus counts in spermatocytes of *Fancj^+/+^* and *Fancj-*deficient animals. Importantly, the current study not only re-examined the effect of the gene-trap on MLH1 focus number, but also utilized two new *Fancj* inactivating mutants – including a full gene deletion – to clarify the meiotic effect of *Fancj* loss. Further, this study marks the first time FANCJ has been visualized during substages of meiotic prophase I, showing a localization pattern highly similar to BRCA1 and TOPBP1. Although we were unable to elucidate a specific mechanism, these data do provide tantalizing clues pertaining to FANCJ’s function in mammalian meiosis.

### FANCJ does not function in meiotic crossover formation

The major finding from this study is that FANCJ does not have a role in CO formation during murine spermatogenesis. Although this conclusion conflicts with our past report^18^, it is strongly supported by the absence of any significant change in either MLH1 foci or chiasmata number in any of the three *Fancj* inactivating mutants analyzed, including the same gene-trap mutant as our previous study and a full gene deletion. It is likely that the differences in strain background might explain the differences between the two reports: In the current study, we used fully backcrossed mouse lines on a C57Bl/6J background, while the previous report used a hybrid C57Bl/6J x 129 mixed background. Further, our current study is strengthened by co-localization and co-immunoprecipitation data that was not available in our previous report. Co-immunofluorescence staining clearly shows that FANCJ does not co-localize with either component of MutLγ (MLH1/MLH3), suggesting FANCJ is absent from the final sites of class I COs. Immunoprecipitation experiments further supported that FANCJ does not interact with MLH1 during meiosis. Additionally, the immunofluorescence signals of MSH4 (a component of MutSγ) and FANCJ appear to be largely independent of each other in late prophase I, though occasional co-localizing foci are observed in early prophase I. This finding is particularly notable as recent reports from our lab have shown correct function of MutSγ might be necessary for all COs^44^. Finally, the timing of FANCJ appearance is inconsistent with a role in class I CO formation, presenting earlier in prophase I than the appearance of CO events. Thus, the conspicuous absence of FANCJ from late MSH4 sites, along with the earlier timing of its localization of chromosome cores suggests that FANCJ function in meiosis is temporally and spatially removed from sites of prospective COs, and likely does not have any direct role in their formation.

### FANCJ functions in meiotic DSB repair

Although FANCJ may not be involved in CO formation, our data suggest a role for FANCJ in early DSB processing events. We find that loss of FANCJ results in persistence of RAD51 on chromosome axes, indicative of persistent DSBs and/or delayed strand invasion events downstream of resection. Recent studies of restriction enzyme-induced DSBs in human cell lines have shown the interaction of FANCJ with CtIP is important for end resection of DSBs to promote HR^51^. End resection is a critical early step in HR, as it produces the ssDNA necessary for RAD51 filament formation and subsequently strand invasion^52^. Importantly, while Nath and Nagaraju found FANCJ deficiency significantly decreased end resection and thereby resulted in a dramatic decrease of RAD51 foci formation, we did not observe any change in RAD51 foci number in zygotene spermatocytes from *Fancj^-/-^*males. This finding suggests that FANCJ is dispensable for correct RAD51 loading, and therefore may not be as necessary for meiotic end resection as it is in somatic contexts. However, *Fancj^-/-^* pachytene spermatocytes showed a significant increase in unrepaired DSBs, as evidenced by aberrant RAD51 and γH2AX autosomal signal. It is tempting to propose that these persistent DSBs are destined to become NCOs, which would suggest FANCJ functions to promote synthesis-dependent strand annealing (SDSA) or some other form of NCO recombination outcome^10,11,53,54^.

In early prophase I, BLM is thought to facilitate branch migration and decatenation of dHJ to produce NCOs^55,56^. Although early involvement of FANCJ with BLM in the regulation of DSB resection and D-loop formation is possible, it remains unknown whether FANCJ associates with BLM in the dissolution of later repair intermediates. On the other hand, the occasional association of FANCJ with MSH4 (which is thought to mark dHJs ^57^, and perhaps earlier DSB repair intermediates) in early and mid zygonema might suggest that FANCJ participates in removal of MutSγ sites that are not destined to become crossovers, or in dHJ processing events after MutSγ removal. However, a direct role for FANCJ in facilitating the unwinding of dHJs alongside BLM remains unclear and somewhat contentious. Prior characterization of BLM in mouse meiosis described localization on synapsed chromosome cores beginning in mid to late zygonema^58^, whilst other studies have described BLM colocalization with RAD51 and DMC1 in leptonema and zygonema^59,60^. Unfortunately, the lack of consistent anti-BLM antibodies makes this difficult to resolve this apparent contradiction at the current time. Our characterization of FANCJ localization primarily to asynapsed regions in late zygonema would thus suggest the two proteins are not likely to co-localize during later stages of prophase I, but could interact functionally prior to dHJ formation. Thus, it remains possible that FANCJ works cooperatively with BLM to achieve D-loop displacement to promote NCO resolution of DSBs via SDSA in early prophase I concurrent with MutSγ removal.

A role for FANCJ in early meiotic NCO formation during DSB repair seems most likely given the prominent localization pattern for FANCJ in early to mid zygonema and what is currently known about FANCJ substrate specificity. FANCJ has been shown previously to unwind forked duplexes and D-loop structures^61^, and *in vitro* studies have demonstrated FANCJ displaces RAD51 in an ATP hydrolysis-dependent manner to suppress RAD51-dependent strand invasion^62^. If FANCJ operates in a similar capacity in meiosis to regulate DNA strand exchange, it could explain our observation of persistent RAD51 foci in *Fancj^-/-^* pachytene spermatocytes. Further support for a role for FANCJ in disrupting D-loops to promote SDSA come from studies of FANCJ deficient human cell lines that reported an increased abundance of aberrant long-tract gene conversions^63^. Interestingly, this suppression of long-tract gene conversions was found to be dependent on the FANCJ-BRCA1 interaction^63^. The function of BRCA1 in regulating DSB resection, strand invasion, and D-loop formation as early steps in SDSA have been described in human cell lines^64–66^. New evidence from *C. elegans* further suggests a role for BRCA1 (BRC-1) in meiotic SDSA/NCO^67^. Although we were unable to confirm a FANCJ-BRCA1 interaction in mouse meiosis by mass spectrometry, the two proteins nearly perfectly co-localize from early zygonema through early pachynema, suggesting their coordinated function.

### Meiotic co-localization of BRCA1, TOPBP1, and FANCJ

Using both a custom antibody against FANCJ and tagging the endogenous protein, our current study provides striking evidence that FANCJ is widely localized as discrete foci along chromosome axes in early meiotic prophase I and as dense accumulations on asynapsed regions in late zygonema and the unsynapsed XY chromosome regions in early pachynema. FANCJ signal strongly co-localized with TOPBP1 and BRCA1 throughout zygonema and early pachynema. BRCA1 forms diverse protein complexes that each have specific molecular functions; the BRCA1-B complex is formed with TOPBP1 and FANCJ and is thought to partially regulate DSB processing, end resection, and S-phase checkpoint signaling^68^. Although we found TOPBP1 and FANCJ to strongly colocalize in early meiotic prophase I, immunoprecipitation experiments suggested they do not interact as we would expect them to if we were detecting the BRCA1-B complex. The TOPBP1-FANCJ interaction is mediated by an S phase-specific phosphorylation on FANCJ at Thr1133^25^, and it is possible FANCJ lacks the requisite phosphorylation to interact with TOPBP1 in meiosis.

TOPBP1 and BRCA1 are well known to promote meiotic silencing of unsynapsed chromatin (MSUC) and MSCI^69,70^. The strong co-localization of FANCJ with TOPBP1 and BRCA1, especially on the sex chromosomes in early pachynema and unsynapsed chromatin in late zygonema, make it tempting to speculate about roles of FANCJ in promoting transcriptional silencing. However, our analysis of *Fancj* mutants has not yet provided any evidence of silencing defects in meiosis. Further, because FANCJ signal on the XY begins to diminish after early pachynema, it is unlikely that FANCJ plays a critical role in promoting or maintaining MSCI. The mechanistic significance of FANCJ colocalization with both TOPBP1 and BRCA1 remains unclear. However, our data are consistent with the possibility that BRCA1 forms different complexes with TOPBP1 and FANCJ that both function in early DSB processing.

### Defining a role for FA proteins in meiosis

Previous studies have reported mice homozygous for the *Fancj* gene-trap disruption showed unaffected G4 DNA metabolism, contrasting with findings in human cell lines that linked loss of FANCJ to G4 DNA-associated genomic instability^31,71^. Such inter-species differences highlight an important caveat in extrapolating mouse data to humans; although we found *Fancj* loss has subtle effect on mouse meiosis, it is possible human patients could experience more severe meiotic phenotypes. However, characterizing meiotic effects of human *Fancj* loss is complicated by high degree of heterogeneity present among clinically identified mutations. For instance, a recent screen of 450 patient-derived mutations found only 12% to cause a loss of function phenotype (defined by heightened sensitivity to ICL-inducing agents)^72^. More broadly, the meiotic effects of FA mutations (as a group) remain poorly understood. A large proportion of FA proteins appear to be present in meiotic prophase. Studies in diverse model organisms have linked FA proteins to roles in meiotic recombination^73–81^, while mouse studies have further implicated several FA proteins with unexpected functions in histone modification and MSCI^82–84^.

Our current report adds to this growing understanding of the complexity of FA protein functions in meiosis. Collectively, these works underscore that FA proteins have idiosyncratic roles in meiosis that cannot be predicted from an understanding of their respective function in somatic DNA repair contexts. Revealing the true role of FA proteins, including FANCJ, may require examining the effects of environmental factors and genetic heterogeneity to uncover compounding effects. Advancing clinical treatments increase life expectancy for patients with FA, there is a growing demand to understand and develop treatments for FA-associated infertility. Elucidating the complex interplay of FA proteins in the unique contexts of meiotic DNA repair and chromatin modification will be critical to these efforts.

### Limitations of study

We encountered significant limitations of our tagged FANCJ and our custom antibody during mass spectrometry. Our custom FANCJ antibody showed numerous non-significant bands in Western blots, and showed non-specific immunofluorescence signal on the XY in late prophase I. We have attempted to validate five putative interactors identified in our initial MS run with this antibody, but all appeared to be non-specific. We have performed protein BLAST searches of our antibody’s antigen sequence, but the only hits with good E-values and more than 50% identity all pertained to FANCJ. Our second MS attempt utilized our FANCJ-HA tag mouse; however, the overall abundance of FANCJ in our elution was too low to make conclusive predictions about putative interactors. Our tag consists of a single HA and 6xHis; although the ability to pull-down FANCJ would be expected to be greater with multiple HA tags, we elected to use a more conservative, single HA out of concern a larger tag would disrupt FANCJ protein stability and function on chromatin. Using the tools currently available, it is not feasible at this time to further pursue FANCJ interactors in mammalian meiosis.

## Materials and Methods

### Generating *Fancj* mutants by CRISPR/Cas9

Production of the *Fancj^GT^* mouseline was previously described.^18^ In association with the Stem Cell and Transgenic Core Facility at Cornell University, we generated three *Fancj* transgenic mouselines using CRISPR/Cas9. Guide RNAs and homology directed repair templates were ordered from IDT and were injected along with Cas9 mRNA into C57BL/6J zygotes.

*Fancj^ΔN^:* Two guide RNAs (5’-GTCTGACTACACCATCGGTG-3’ and 5’- AAGCTAGTCGTGCATCTACA-3’, targeting chromosome 11: 86198931-86198950 and 86187036-86187055, respectively) were used to create a 11,914 bp deletion beginning ∼33 bp downstream of the start codon. The deleted region was predicted to encompass the coding sequences for the MLH1 interaction site, the nuclear localization domain, and much of the N-terminal helicase ATP-binding domain.182 embryos were injected, and 125 2-cell embryos were surgically implanted into 4 recipient female mice. A total of 21 founders were born, and PCR screening identified 10 founders carrying the deletion.

*Fancj^-^:* Two guide RNAs (5’-CAGCTCTATTGCTAGGTGTC-3’ and 5’- CGCCAGCATCAGCTAATCCC-3’, targeting chromosome 11: 86199222-86199241 and 86061270-86061289, respectively) were used to delete the entire coding sequence of the *Fancj* gene beginning 277 bp upstream of the start codon and ending 159 bp downstream of the stop codon. Donor DNA consisting of 78 nucleotides of homology to the 5’ and 3’ UTRs flanking six stop codons was used to replace the entirety of the *Fancj* coding sequence via Homology directed repair. 189 embryos were injected, and 117 2-cell embryos were surgically implanted into 4 recipient female mice. A total of 9 founders were born, and PCR screening identified 1 founder carrying the deletion.

*Fancj^HA^:* CRISP/Cas9 insertion of a 6xHis-HA epitope tag sequences to the endogenous *Fancj* locus was done using one guide RNA (5’-TGGGCAGAAGTGAAAGTCGA-3’ targeting chromosome 11: 86061422-86061441) and donor DNA consisting of 80 nucleotide homology upstream of the guide sequence, 6xHis and HA tag sequences, a mutated PAM sequence, stop codon and 80 nucleotides homology downstream of the stop codon. 182 embryos were injected, and 140 2-cell embryos were surgically implanted into 4 recipient female mice. A total of 15 founders were born, and PCR screening identified 4 founders carrying the insertion of the 6xHis-HA tag immediately prior to the endogenous stop codon.

### Care and use of experimental animals

All moue strains were backcrossed at least two generations and maintained on a C57BL/6J background (Jackson Laboratory). Mice were housed in polysulfone cages on ventilated racks (Allentown PC75JHT). Cages contained Bed-o’-cobs ¼” bedding (Scotts 330BB), Bio-Serv standard hut (Scotts K3352), and Crinkle Nest Kraft (Scotts CNK), and a 1” x 1” nestlet (Scotts NES7200) for enrichment. Mice were maintained under strictly controlled conditions of constant temperature, 12-hour light/dark cycles, and provided food (Envigo Bioproducts Inc Rodent diet 7012) and water *ad libitum*. All mice in this study were killed between 6 and 18 wks. of age. Animal handling and procedures were performed following approval by the Cornell Institutional Animal Care and Use Committee under protocol 2004-0063.

### Genotyping mice, cDNA preparation and analysis

Genomic DNA was extracted from mouse ear snips at weaning. All genotyping was done with 10 μM primers and using OneTaq 2X master mix (NEB M0482) according to the manufacturer’s recommendations. To validate genotyping, samples were run on 1.5% agarose, bands were cut and extracted using a QIAquick Gel Extraction Kit (Qiagen 28704) and were submitted to the Cornell Biotechnology Resource Center for Sanger sequencing.

*Fancj^GT^*: Mice were genotyped via two separate reactions using a common forward primer paired with one of two reverse primers:

> EX5 1-S (common forward): 5’-TGCCAAGAAACAGGCATCTATAC-3’
>
> EX5 1-1000-AS (wildtype reverse): 5’-ATGACCTCTTCTGATCTCTGCTG-3’
>
> EN2 Intr (gene trap reverse): 5’-GGCCTGCTCAAACCTGAACC-3’

The wildtype allele generated a 972 bp band while the gene trap allele generated a 380 bp band. PCR conditions were the same for both wildtype and gene trap reactions and included 55°C annealing with a 1 min extension for 35 cycles.

*Fancj^ΔN^:* Mice were genotyped via two separate reactions using a common forward primer paired with one of two reverse primers:

> Fancj-A (common forward): 5’-TGGCTATGCCAGGTAAATGC-3’
>
> Fancj-B (wildtype reverse): 5’-AGGCCTCAGGCACACATAAG-3’
>
> Fancj-D (*ΔN* reverse): 5’-CGCCCAACAAACAAGTAGAC-3’

The wildtype allele generated a 731 bp band while the ΔN allele generated a 727 bp band. PCR conditions were the same for both wildtype and ΔN reactions and included 50°C annealing with a 1 min extension for 35 cycles.

*Fancj^-^*: Full null mice were genotyped using the following primers: FancJ-F (5’- TACCTGAGGACATGGAGTTCTA-3’) and FancJ-R (5’-CCAGGCAACTAAGTATCAGAGG-3’).

The *Fancj^-^* allele generated a 407 bp band, while wildtype alleles were detected using the Fancj-A and Fancj-B primers above. PCR conditions for the full null allele included 53°C annealing with a 1 min extension for 35 cycles.

*Fancj^HA^*: The epitope tagged *Fancj* allele was genotyped using the following primers: FancJ-tag_FWD (5’-GTCGCCAGGAACTGAAGAAG-3’) and FancJ-tag_REV (5’- TGCTCTCTTCCACTGGTGTG-3’). The wildtype allele generated a 277 bp band while the tagged allele generated a 322 bp band. PCR conditions included 46.6°C annealing with a 1 min extension for 30 cycles. Confirmation of the epitope tag insertion was further validated using the same PCR condition but with an alternative reverse primer, FancJ-intag_REV (5’- GTAATCTGGAACATCGTATG-3’), corresponding to part of the HA-tag coding sequence. The tagged allele generated a 259 bp band while the wildtype allele did not generate any band.

Mouse *Fancj* transcriptomic analysis was done using RNA extracted from thymus or testis using TRIzol reagent (Invitrogen 15596026). cDNA was produced using the SuperScript III Reverse Transcriptase kit (Invitrogen 18080093). Assessment of the transcription of *Fancj* in mice was assessed by PCR using the following primers: FancJcDNA_F (5’- TGTCCTCAGTGTTGTCTGAC-3’), which corresponds to sequence in exon 2, and FancJcDNA_R (5’-GAAATTCCCCACCACTTCA-3’), which corresponds to sequence in exon 7. The wildtype transcript generated an 884 bp band while the *ΔN* transcript generated a 373 bp band and the full null produced no transcript and thus no band. PCR conditions included a 66°C annealing with a 1 min extension for 35 cycles.

### HSPC quantification by flow cytometry

Bone marrow cells were isolated from femurs, tibiae and iliac crests with 10 ml ice cold PBS supplemented with 2 % fetal calf serum (FCS) and strained through 70 μm cell strainers. Red cells were lysed by resuspending the cells in 10 ml red cell lysis buffer (#46232, Cell Signaling Technology) for 10 minutes at room temperature. After centrifugation, the cell pellet was resuspended in PBS supplemented with 2 % FCS and nucleated cells were counted in methylene blue with 3 % acetic acid on a Vi-Cell XR cell viability counter (Beckman Coulter). 10 × 10^6^ bone marrow cells were resuspended in 200 μl of PBS supplemented with 2 % FCS containing the following antibody solution: FITC-conjugated lineage cocktail with antibodies against CD4 (clone H129.19, BD Pharmingen), CD3e (clone 145-2C11, eBioscience), Ly-6G/Gr-1 (clone RB6-8C5, eBioscience), CD11b/Mac-1 (clone M1/70, BD Pharmingen), CD45R/ B220 (clone RA3-6B2, BD Pharmingen), FcεR1α (clone MAR-1, eBioscience), CD8a (clone 53-6.7, BD Pharmingen), CD11c (clone N418, eBioscience), TER-119 (clone Ter119, BD Pharmingen); c-Kit (PerCP-Cy5.5, clone 2B8, eBioscience), Sca-1 (PE-Cy7, clone D7, eBioscience), Flt3 (PE, clone A2F10, eBioscience), CD34 (eFluor660, clone RAM34, eBioscience), CD16/32 (BV421, clone 93, BioLegend) and Il-7R⍺ (BV605, clone A7R34, BioLegend). The cells were incubated in the antibody solution for 90 minutes at room temperature, washed in 3 mls of PBS supplemented with 2 % FCS, centrifuged, resuspended in 800 μL of PBS supplemented with 2 % FCS and analyzed on Attune NxT Flow Cytometer (Thermo Fisher Scientific).

### Sperm count

For each adult mouse analyzed, both caudal epididymides were removed and placed in 1 mL of prewarmed DMEM containing 4% bovine serum albumin (Sigma-Aldrich). Sperm were extruded from each epididymis using tweezers and subsequently incubated at 37°C/5% CO2 for 20 minutes. A 20 μl aliquot of the sperm suspension was resuspended in 480 μl 10% formalin, and the sperm cells counted using a hemocytometer.

### Generation of antibody against FANCJ

In association with Thermo Fisher Scientific, we made a custom rabbit polyclonal antibody against the murine FANCJ C-terminus (antigen sequence: DDSECFTPELFDPVDTNEE corresponding to residues 1124-1142). Host animals were immunized and received boosted immunization before terminal bleed. Serum from terminal bleeds was affinity purified.

### Spermatocyte chromosome preparations and immunofluorescence staining

Spermatocyte prophase I chromosome spreads were made according to the method developed by Peters and colleagues^85^. Briefly, testes were cleaned and detunicated in PBS and incubated in hypotonic extraction buffer (30 mM tris; 50 mM sucrose; 17 mM sodium citrate; 5 mM EDTA; 5 mM PMSF; 2.5 mM DTT; pH 8.2-8.4) for 20 min on ice. Several seminiferous tubules from each testis were separated and macerated in 500 mM sucrose at room temperature. Cell suspensions were spread on slides coated in 1% paraformaldehyde (pH 9.2-9.3) with 0.15% Triton X. Slides were incubated in a room temperature humid chamber for 2 hours and washed with 0.4% Photo-flo 200 solution (Kodak Professional) for 2 min. Slides were air-dried for 30 min then either stored at −80°C or immediately stained. For staining, slides were washed in PBS with 0.4% Photo-flo for 10 minutes, followed by 10 minutes in PBS with 0.1% Triton X, and finally for 10 minutes in 10% antibody dilution buffer (3% bovine serum albumin, 10% normal goat serum, 0.0125% Triton X, and PBS). Primary antibodies were diluted in 100% antibody dilution buffer, applied to slides, covered with a parafilm coverslip, and incubated in a room temperature humid chamber overnight. Slides were washed in PBS with 0.4% Photo-flo for 10 minutes, followed by 10 minutes in PBS with 0.1% Triton X, and finally for 10 minutes in 10% antibody dilution buffer. Secondary antibodies were diluted and applied to slides in a similar fashion as the primary antibodies. Slides were incubated in a room temperature humid chamber for 2 hours, washed 3x 5 minutes in PBS with 0.4% Photo-flo, the 1x 5 minutes in Milli-Q water with 0.4% Photo-flo. Finally, slides were mounted using 30 μl ProLong Diamond antifade with DAPI (Invitrogen P36962) and glass coverslips were applied. Slides were stored at 4°C prior to analysis.

Antibodies and dilutions used in this study include: MLH1 (mouse) mAb 1:50 (BD Biosciences 550838); MLH3 (guinea pig) pAb 1:500 (custom made with Thermo-Fisher); RAD51 (rabbit) pAb 1:500 (Millipore PC130); γH2AX (mouse) mAb 1:10,000 (Millipore 05-636); MSH4 (rabbit) pAb 1:100 (ABclonal A8556); FANCJ (rabbit) pAb 1:100 (custom made with Thermo-Fisher; detailed above); HA-Tag (C29F4) (rabbit) mAb 1:100 (Cell Signaling Technology 3724); HA-Tag (6E2) (mouse) mAb 1:100 (Cell Signaling Technology 2367); SYCP3 (mouse) mAb 1:1,000 (Abcam ab97672); SYCP3 (rabbit) 1:10,000 (custom made^12^); BRCA1 (rabbit) 1:100 (from Raimundo Freire); and TOPBP1 (rabbit) serum antibody 1:100 (from Raimundo Freire). All secondary antibodies were from Jackson ImmunoResearch and were diluted 1:1000, including: AffiniPure F(ab’)2 fragment goat anti-mouse IgG Fc fragment specific conjugated to AF488 (115-546-008) or AF594 (115-586-008); anti-rabbit IgG Fc fragment specific conjugated to AF488 (111-546-046) or AF594 (111-586-046); and AffiniPure F(ab’)2 fragment goat anti-guinea pig IgG conjugated to AF647 (106-606-003).

### Diakinesis spermatocyte preparations for observation of chiasmata

Spermatocyte diakinesis chromosome spreads were prepared according to methods developed by Evans and colleagues^86^. Briefly, testes were detunicated and seminiferous tubules were macerated in 2.2% sodium citrate. Tubules were further dissociated by pipetting and larger fragments of tubules were left to settle for 15 minutes. The supernatant cell suspension was collected and centrifuged for 5 minutes at 600xg. The resulting cell pellet was resuspended in 4 mL 1.1% sodium citrate and incubated at room temperature for 15 minutes. The cell suspension was centrifuged for 5 min at 600xg and the sodium citrate supernatant was discarded. The cell pellet was vigorously vortexed into suspension as fixative (3:1 methanol:glacial acetic acid) was slowly added to the cells. Following 2 additional centrifugations and resuspension in fixative, cells were dropped on slides and air dried. Slides were stained for 6 minutes with 4% Giemsa (Fisher), washed three times for 3 minutes each with ddH2O, dried, and mounted with permount.

### Image acquisition

For immunofluorescence experiments, slides were imaged on a Zeiss Axio Imager epifluorescence microscope at 63x magnification. DAPI, AF488, AF594, and AF647 were imaged sequentially using standardized exposure times for each antibody condition and processed using Zeiss Zen Blue version 3.0 (Carl Zeiss AG, Oberkochen, Germany). Images were adjusted in ImageJ (National Institutes of Health, USA) to standardize background across all images. Diakinesis cells stained with Giemsa were imaged on a Zeiss Axio Imager epifluorescence microscope at 40x magnification using brightfield. Images were processed using Zeiss Zen Blue version 3.0 (Carl Zeiss AG, Oberkochen, Germany). In most cases, SYCP3 was used to sub-stage cells, with the extent of XY synapsis used to substage pachytene into early, mid and late. For FANCJ colocalization studies with TOPBP1, MSH4, MLH1, and BRCA1, cells were sub-staged by DAPI. Leptonema/zygonema were defined by 21-40 centromeres (DAPI-dense spots) without a visible sex body; pachynema having 21 centromeres and a defined sex body; and diplonema having more than 21 centromeres and a defined sex body. Zygotene cells could be further categorized by early/mid and late using FANCJ staining.

### Germ cell enrichment and protein extraction

Enriched populations or total germ cells were extracted from testes of two age-matched mice per replicate for immunoprecipitation experiments. Testes were washed and detunicated in PBS. Seminiferous tubules were macerated in PBS and further dissociated by pipetting. The resulting cell suspension was filtered twice, first through a 70 μm cell strainer then a 40 μm strainer. Filtered cell suspensions underwent three rounds of centrifugation (5 minutes at 600xg) and subsequent resuspension in PBS. Following the final centrifugation, the cell pellet was resuspended in 750 μl Tris lysis buffer (50 mM Tris, 0.2% NP-40, 150 mM NaCl, 5 mM EDTA, 5.7 mM PMSF, and 1 x Roche cOmplete). For whole organ samples (either whole detunicated testis or thymus), the tissue was placed directly in 1 mL Tris lysis buffer. Samples were sonicated 2x for 12 seconds at 23% amplitude in cycles of 0.4 seconds on and 0.2 seconds off. Samples were then centrifuged for 20 minutes at 15,000xg at 4°C, and the lysate was either used immediately for downstream applications or stored at −80°C.

### Immunoprecipitation

Sample lysates were precleared by incubation with Dynabeads protein A (Invitrogen 10001D) for 1 hour at 4°C on a nutator. Beads were then pelleted with a magnet and the lysate supernatant was transferred to be incubated with either Dynabeads and 10 μg (rabbit)FANCJ antibody or with 10-25 μl (rabbit)HA antibody covalently bound to magnetic beads (Cell Signaling Technology 11846S). Lysate, bead, and antibody mixtures were incubated overnight at 4°C on a nutator. Samples were then subjected to 4-8 cycles of pelleting on magnet, discarding of the supernatant flow through, and washing in Tris lysis buffer. Following the final wash, beads were resuspended in an elution buffer containing 100mM Tris, 1% SDS, and 10mM DTT. Samples were eluted at 65°C for 15 minutes. Elution was collected and either used immediately for downstream applications or stored at −80°C.

### SDS-PAGE and Western blotting

Cell and tissue lysate samples were separated by SDS-PAGE on bis-acrylamide gels varying in percentage from 8% - 12.5% and transferred to methanol activated PVDF membranes using a Biorad Mini Trans-Blot Cel. Membranes were incubated in EveryBlot Blocking Buffer (Biorad 12010020) for 10 minutes at room temperature while rotating at 60 rpm. Primary antibodies were diluted in EveryBlot Blocking Buffer and incubated with the membrane overnight at 4°C while rocking. Membranes were washed 3x for 15 minutes in 1X TBST and subsequently incubated with secondary antibodies in EveryBlot Blocking Buffer for 1 hour at room temperature. Membranes were washed 3x for 15 minutes in 1X TBST and finally developed using the ECL reagent and imaged using a Biorad ChemiDoc XRS+ imaging system.

Antibodies and dilutions used in this study include: MLH1 (mouse) mAb 1:500 (BD Biosciences 550838), FANCJ (rabbit) pAb 1:1000 (custom made with Thermo-Fisher); HA-Tag (C29F4) (rabbit) mAb 1:1000 (Cell Signaling Technology 3724); HA-Tag (6E2) (mouse) mAb 1:1000 (Cell Signaling Technology 2367); TOPBP1 (rabbit) serum antibody 1:1000 (from Raimundo Freire); ATRIP (rabbit) pAb 1:1000 (Thermo-Fisher PA1519); TRIM28 (KIP1) (rabbit) mAb 1:1000; USP4 (mouse) mAb 1:1000 (Proteintech 66822-1-lg); RUVBL1 (rabbit) pAb 1:1000 (Proteintech 10210-2-AP); and ADAD2 (rabbit) 1:2000 (gift from Elizabeth Snyder). Secondary antibodies used included: Pierce goat anti-mouse IgG (H+L) peroxidase conjugated (Thermo 31430) and Pierce goat anti-rabbit IgG (H+L) peroxidase conjugated (Thermo 31460), each used at 1:5000.

### Label-free mass spectrometry and protein identification

Initial label-free mass spectrometry analysis of anti-FANCJ IP from whole testis lysate was performed by the Cornell University Proteomics and Metabolomics Core Facility as previously described^87^. In brief, a nanoLC-MS/MS analysis was performed using an Orbitrap Fusion (Thermo-Fisher Scientific, San Jose, CA) mass spectrometer equipped with a nanospray Flex Ion Source using high energy collision dissociation (HCD) coupled with UltiMate3000 RSLCnano (Dionex, Sunnyvale, CA). Reconstituted samples were injected onto a PepMap C-18 RP nano trap column (3 μm, 100 μm x 20 μm, Dionex) with nanoViper fittings at a 20 μL/min flow rate for on-line desalting, and subsequently separated on a PepMap C-18 RP nano column (3 μm, 75 μm x 25 μm) and eluted in a 120 min gradient of 5-35% acetonitrile (CAN_ in 0.1% formic acid at 200 nL/min. The Orbitrap Fusion was operated in a data-dependent acquisition mode using FT mass analyzer for one survey scan followed by three “Top Speed” data-dependent CID ion trap MS/MS scans with normalized collision energy of 30%. A dynamic exclusion window of 45 seconds was specified. Data was acquired using Xcalibur 3.0 operation software and Orbitrap Fusion 2.0 (Thermo-Fisher Scientific).

Raw spectra for each sample and replicate were processed and proteome databases searched using Proteome Discoverer 2.5 (PD 2.5, Thermo-Fisher Scientific) with the Sequest HT search engine. Search settings included: two missed cleavage sites by trypsin allowed with fixed carboxamidomethyl modification of cysteine and variable modifications of methionine oxidation and deamidation of asparagine and glutamine. Identified peptides were filtered for a maximum of 1% false discovery rate (FDR) using the Percolator algorithm in PD2.5. Relative label-free quantification was done in PD 2.5 to calculate protein abundances. The number of peptide spectrum matches (PSMs) were summed to represent protein abundance. Abundance ratios were calculated based on pairwise ratios using the median calculated among replicates.

### Sample preparation for mass spectrometry and TMT labeling

Proteins eluted from anti-HA beads were processed for mass spectrometry analysis following a previously described protocol that includes reduction, alkylation, precipitation, and digestion by trypsin over-night ^88^. Peptides were then desalted, dried^89^ and labelled with TMT following a protocol described previously^90^. Briefly, the dried peptides were resuspended in 50 mM HEPES, and labeled with 100 µg of TMT sixplex Isobaric Label Reagents (Thermo Scientific) using 3 TMT channels for each sample (*Wild-type* and *FANCJ-HA*). TMT-labeling reactions were done at room temperature for 1h and quenched with 5% hydroxylamine for 15 minutes. After quenching, TMT-labeled peptides from all six channels were pooled, acidified with 0.1% TFA, and desalted using a SPE 1cc C18 cartridge (Sep-Pak C18 cc vac cartridge, 50 mg Sorbent, WAT054955, Waters). Bound TMT-labeled peptides were eluted with 80% acetonitrile, 0.1% acetic acid in water before being dried.

### Mass spectrometric analysis of TMT-labeled peptides

TMT-labeled peptides were analyzed by mass spectrometry as described previously^88,90^. Peptides were pre-fractionated using off-line HILIC chromatography and fractions were analyzed by LC-MS/MS using an UltiMate 3000 RSLC nano chromatographic system coupled to a Q-Exactive HF mass spectrometer (Thermo Fisher Scientific). Reverse-phase capillary chromatography was performed using a 35cm-long 100 µm inner diameter column packed in-house with 3 µm C18 reversed-phase resin (Reprosil Pur C18AQ 3 μm). The Q-Exactive HF was operated in data-dependent mode with survey scans acquired in the Orbitrap mass analyzer over the range of 380–1800 m/z with a mass resolution of 120,000. MS/MS spectra were performed after selecting the top 7 most abundant +2, +3, or +4 ions and a precursor isolation window of 0.7 m/z. Selected ions were fragmented by Higher-energy Collisional Dissociation (HCD) with normalized collision energies of 28, with fragment mass spectra acquired in the Orbitrap mass analyzer with a monitored first mass of 100 m/z, mass resolution of 15,000, AGC target set to 1 × 10^5^, and maximum injection time set to 100 ms. A dynamic exclusion window of 30 seconds was specified.

### Mass spectrometry data analysis

The Trans Proteomic Pipeline (TPP) version 6.0.0 was used for peptide identification and quantification as described previously^88^. Briefly, MS data were converted to mzXML using msConvert. The spectral data files were searched using the Comet search engine (v2021 rev 1)^91^. The validation of peptide identifications was done using PeptideProphet, and TMT channel quantification was performed using Libra. The Results from Libra were exported as tab-delimited files for further processing via R scripts^90^.

### Mammalian Cell Culture

The mouse Pre-B cells was a gift from Barry Sleckman and were grown at 37⁰C/5%CO_2_ in Dulbecco’s modified Eagle medium (DMEM) supplemented with 10% Bovine Calf Serum (BCS), 1% non-essential amino acids, 0.0004% beta mercaptoethanol and penicillin/streptomycin solution (100 U/ml).

### Ionizing Radiation

The mouse Pre-B cells were exposed to 20gy of gamma radiation (0.89 Gy/min) from a Cs-137 source and were harvested 90 minutes after the treatment.

### Quantification and statistical analyses

MLH1 and RAD51 foci that were at least 50% localized to SYCP3 were counted. MLH1 foci were quantified from 25-30 cells per animal, with the following sample sizes foreach genotype: four *Fancj^+/+^* animals (119 cells); two *Fancj^+/GT^* animals (57 cells); seven *Fancj^GT/GT^* animals (204 cells); one *Fancj^+/ΔN^* animal (27 cells); five *Fancj^ΔΝ/ΔΝ^* animals (144 cells); two *Fancj^+/-^* animals (60 cells); and three *Fancj^-/-^* animals (90 cells). Among group differences in mean MLH1 foci counts were analyzed by one-way ANOVA; statistically significant differences (p < 0.05) between each mutant genotype’s mean and *Fancj^+/+^* were determined by Dunnett’s multiple comparison test. RAD51 foci were counted only from *Fancj^+/+^* and *Fancj^-/-^* genotypes, and a minimum of 20 cells per animal per stage. Two animals (45 cells total) per genotype were analyzed for zygotene RAD51, and three animals were analyzed for pachytene RAD51 (73 and 70 cells total for *Fancj^+/+^* and *Fancj^-/-^*, respectively). Statistical differences between groups were determined by Welch’s two-tailed t-test. Chiasmata were counted for 25-30 cells per animal for three *Fancj^+/+^* (85 cells total) and three *Fancj^-/-^* animals (90 cells total). Significant differences between groups was determined by Welch’s two-tailed t-test. For each animal, each testis was weighed independently. The mean mass of both testes was divided by the individual male’s body mass to calculate the proportion of testis mass: body mass. This measurement was made for thirteen *Fancj^+/+^*, five *Fancj^+/GT^*, eight *Fancj^GT/GT^*, four *Fancj^+/ΔΝ^*, nine *Fancj^ΔΝ/ΔΝ^*, four *Fancj^+/-^*, and ten *Fancj^-/-^* males. Among group differences in mean testis mass: body mass was analyzed by one-way ANOVA; statistically significant differences (p < 0.05) between each mutant genotype’s mean and the WT were determined by Dunnett’s multiple comparison test. Sperm counts were collected for eleven *Fancj^+/+^*, five *Fancj^+/GT^*, nine *Fancj^GT/GT^*, four *Fancj^+/ΔΝ^*, nine *Fancj^ΔΝ/ΔΝ^*, four *Fancj^+/-^*, and seven *Fancj^-/-^*males. Among group differences in mean sperm counts were analyzed by one-way ANOVA; statistically significant differences (p < 0.05) between each mutant genotype’s mean and the WT were determined by Dunnett’s multiple comparison test. Statistical differences between *Fancj^+/+^*and *Fancj^-/-^* HSPCs and myeloid progenitor cells were determined by two-tailed Mann-Whitney U test.

## Acknowledgements

We are grateful to Eileen Shu for assistance with mouse husbandry and overall lab management, and the entire Cohen lab for feedback and advice during the course of this research. We thank Barry Sleckman for the pre-B cells; Raimundo Freire for the BRCA1 and TOPBP1 antibodies; Elizabeth Snyder for the ADAD2 antibodies; Rob Munroe and Chris Abratte of the Cornell Transgenic Mouse Core Facility for the generation of new mouse lines; and the Cornell Center for Animal Resources and Education for providing animal care and veterinary services to research animals. We also thank the Proteomics and Metabolomics Facility of Cornell University for providing label-free mass spectrometry data, funding for the Orbitrap Fusion mass spectrometer was provided to Dr. Sheng Zhang from NIH SIG (1S10 OD017992-01). Work described in this manuscript was supported by funding to P.E.C. from the Eunice Kennedy Shriver National Institute for Child Health and Development (HD097897) and funding to M.B.S. from the Eunice Kennedy Shriver National Institute of Child Health and Human Development (HD095296).

