## Supplementary Figures for "The DNA helicase FANCJ (BRIP1) functions in Double Strand Break repair processing, but not crossover formation during Prophase I of meiosis in male mice"

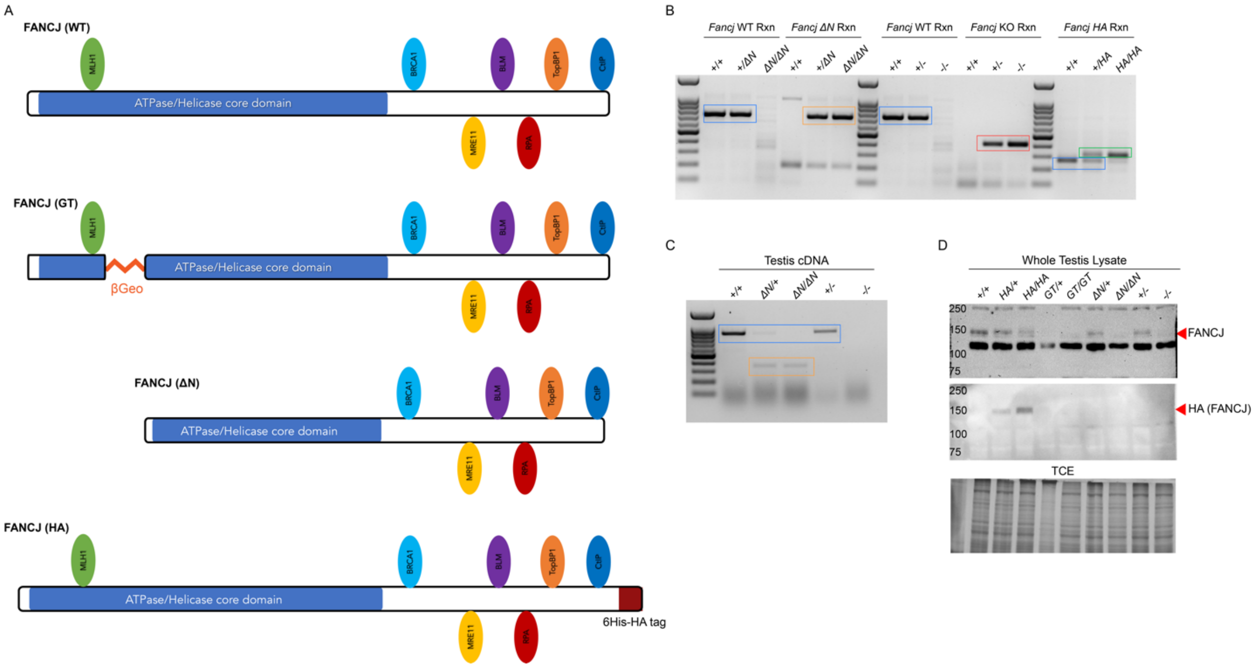


**Figure S1 Validation of *Fancj* mutants and antibodies.** A) Diagram detailing the wildtype, gene-trap, ΔN, and epitope-tagged FANCJ proteins with their predicted protein interaction sites (The full deletion is excluded as it does not make a protein). The FANCJ-ΔN protein in missing the N-terminal region of the ATPase/Helicase core domain and the MLH1 interaction site, while the epitotpe-tagged protein has a 6xHis-HA dual tag at the C-terminus. B) PCR genotyping of *Fancj*$^{\Delta N}$, *Fancj*$^{-}$, and *FancJ*$^{HA}$ mutants. PCR product bands are boxed in colors pertaining to allele: *Fancj*$^{+}$ (blue), *Fancj*$^{\Delta N}$ (orange), *Fancj*$^{-}$ (red), and *Fancj*$^{HA}$ (green). C) PCR of testis cDNA from *Fancj*$^{+/+}$, *Fancj*$^{+/\Delta N}$, *Fancj*$^{\Delta N/\Delta N}$, *Fancj*$^{+/-}$, and *Fancj*$^{-/-}$ mice. PCR product bands are boxed in colors pertaining to allele coding sequence: *Fancj*$^{+}$ (blue), *Fancj*$^{\Delta N}$ (orange). As expected, no band was detected for *Fancj*$^{-}$, indicting no transcript was made. D) Western blots using whole testis lysate from each *Fancj* genotype: *Fancj*$^{+/+}$, *Fancj*$^{HA/+}$, *Fancj*$^{HA/HA}$, *Fancj*$^{GT/+}$, *Fancj*$^{GT/GT}$, *Fancj*$^{\Delta N/+}$, *Fancj*$^{\Delta N/\Delta N}$, *Fancj*$^{+/-}$, and *Fancj*$^{-/-}$. Lysates were run on an 8% bis-acrylamide, 0.5% TCE gel. Top depicts the membrane blotted with rabbit antibody to FANCJ; middle is the same membrane after being stripped and reblotted with mouse antibody to HA; and bottom is the TCE loading control. Red arrows indicate FANCJ and HA bands (expected size: 131 kDa but detected at 150 kDa).

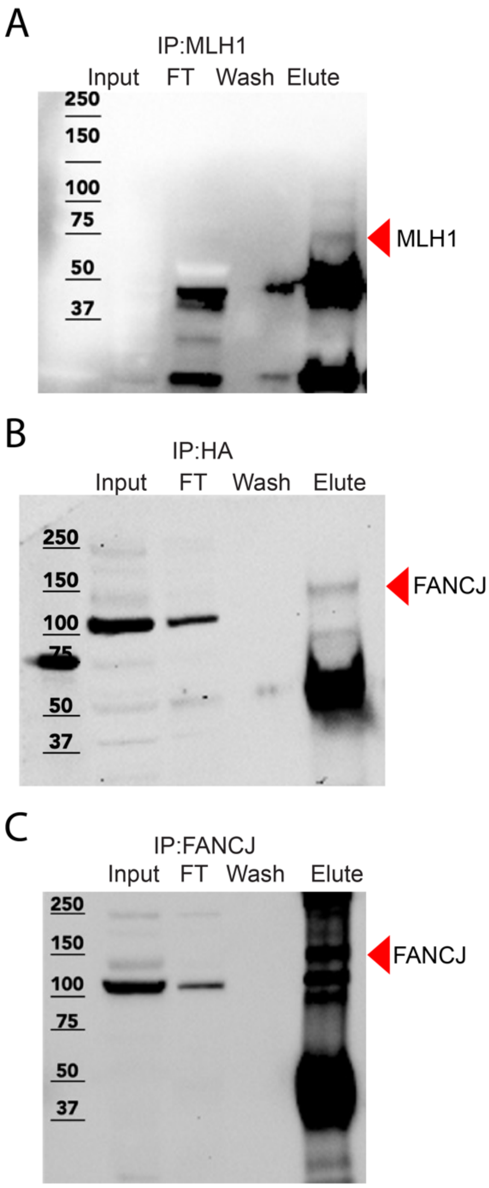


**Figure S2 MLH1 and FANCJ detected in their respective IP-WBs.** Reciprocal IP-WBs from those shown in Fig 5. A) Western blot of MLH1-IP using whole testis lysate input. Sample was run on an 8% SDS-PAGE gel, and membrane blotted with antibody against MLH1. Black arrow points to band corresponding to MLH1 in the elution. B) Western blot of HA-IP using *Fancj*$^{HA/HA}$ whole testis lysate input. Sample was run on an 8% SDS-PAGE gel, and membrane blotted with antibody against FANCJ. Black arrow highlighting FANCJ bands in elution. C) Western blot of FANCJ-IP using whole testis lysate input. Sample was run on an 8% bis-acrylamide gel, and membrane blotted with antibody against FANCJ. Black arrow highlighting FANCJ bands.

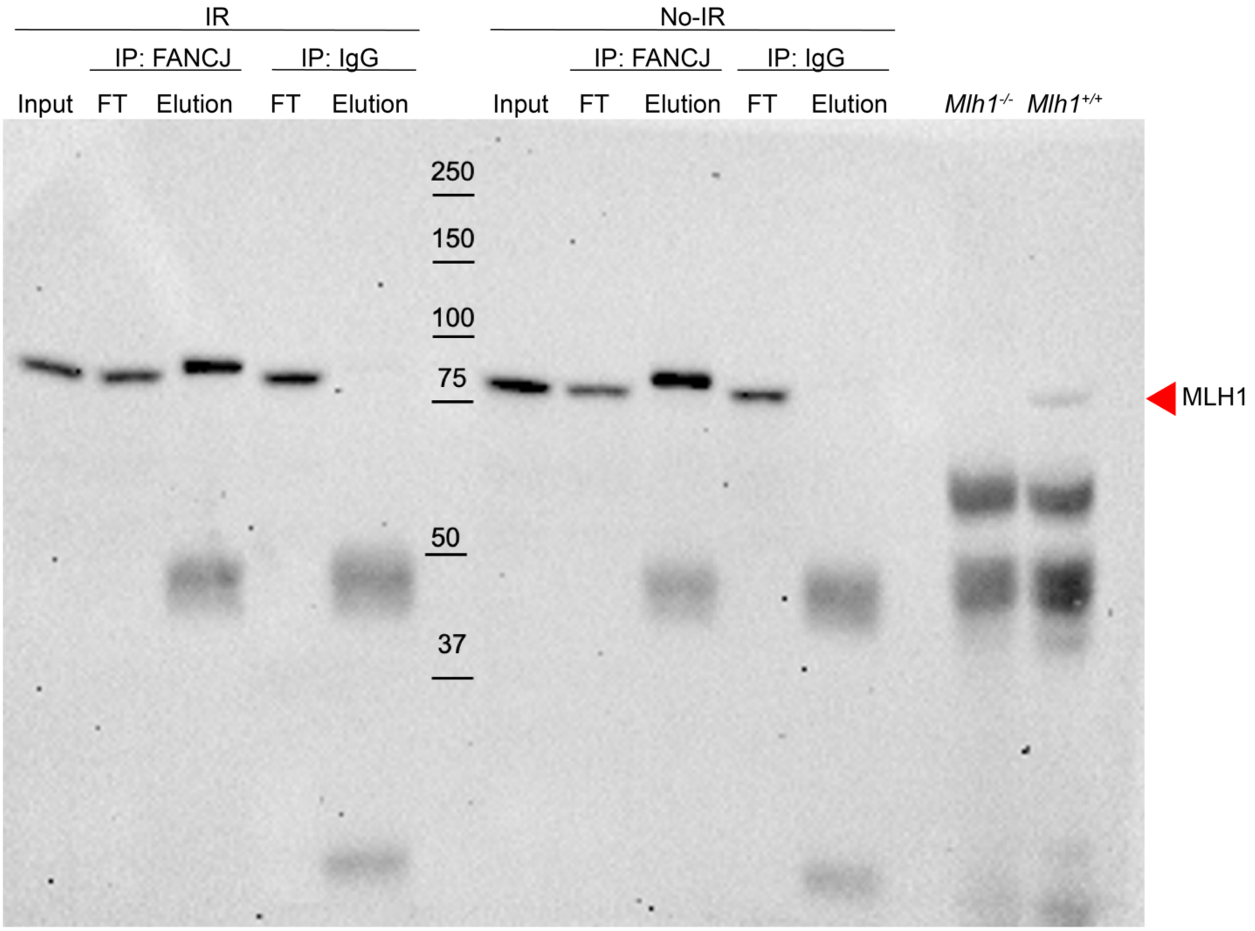


**Figure S3 FANCJ interacts with MLH1 in pre-B cells**. Lysate from irradiated (IR) and untreated (No-IR) pre-B cells was immunoprecipitated using antibody against FANCJ. The input lysates, flow throughs, and elutions were run on an 8% bis-acrylamide gel. The membrane was blotted with an antibody against MLH1. Red arrow points to MLH1 bands. Lysate from *Mlh1*$^{+/+}$ and *Mlh1*$^{-/-}$ mouse whole testis lysates was also run as a positive and negative control.

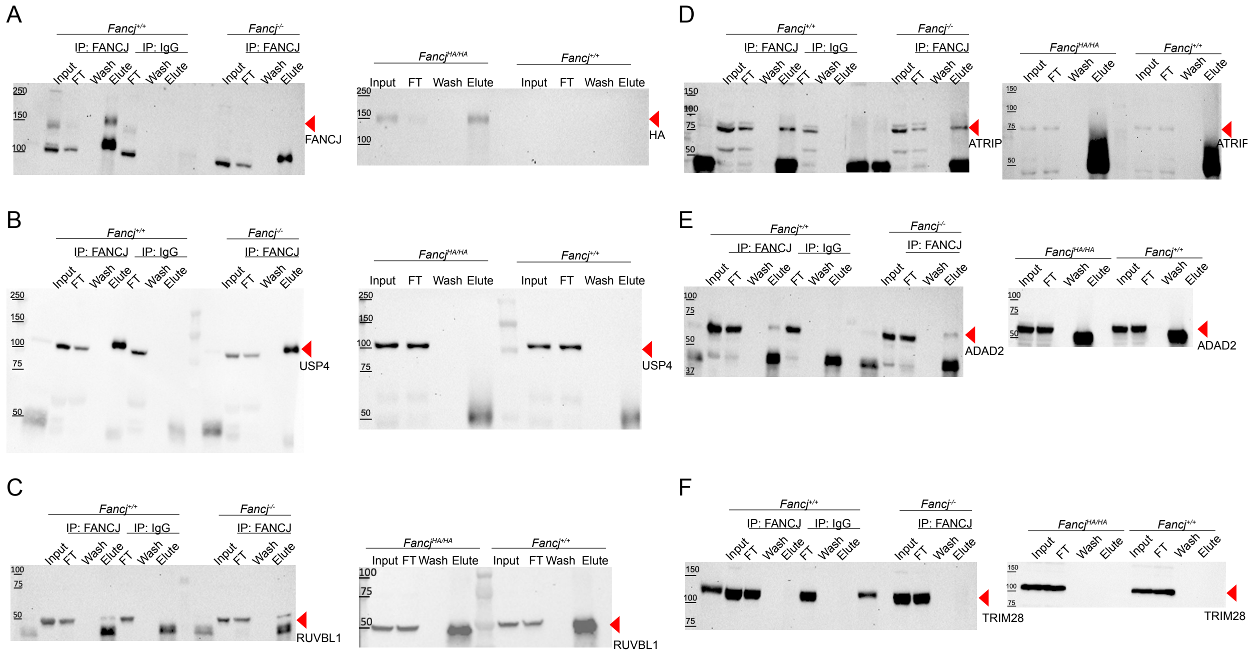


**Figure S4 Putative interactors found in initial FANCJ IP-MS experiments were non-specific.** Western blots of immunoprecipitation experiments to validate initial FANCJIP-MS experiments. Each set of validations consisted of 5 separate IPs: FANCJ-IP and IgG-IP from *Fancj*$^{+/+}$ whole testis lysate; FANCJ-IP from *Fancj*$^{-/-}$ whole testis lysate; and HA-IP from *Fancj*$^{HA/HA}$ and *Fancj*$^{+/+}$ whole testis lysates. Membranes were blotted with antibodies to: A) FANCJ and HA (to validate the IPs), B) USP4, C) RUVBL1, D) ATRIP, E) ADAD2, and F) TRIM28.

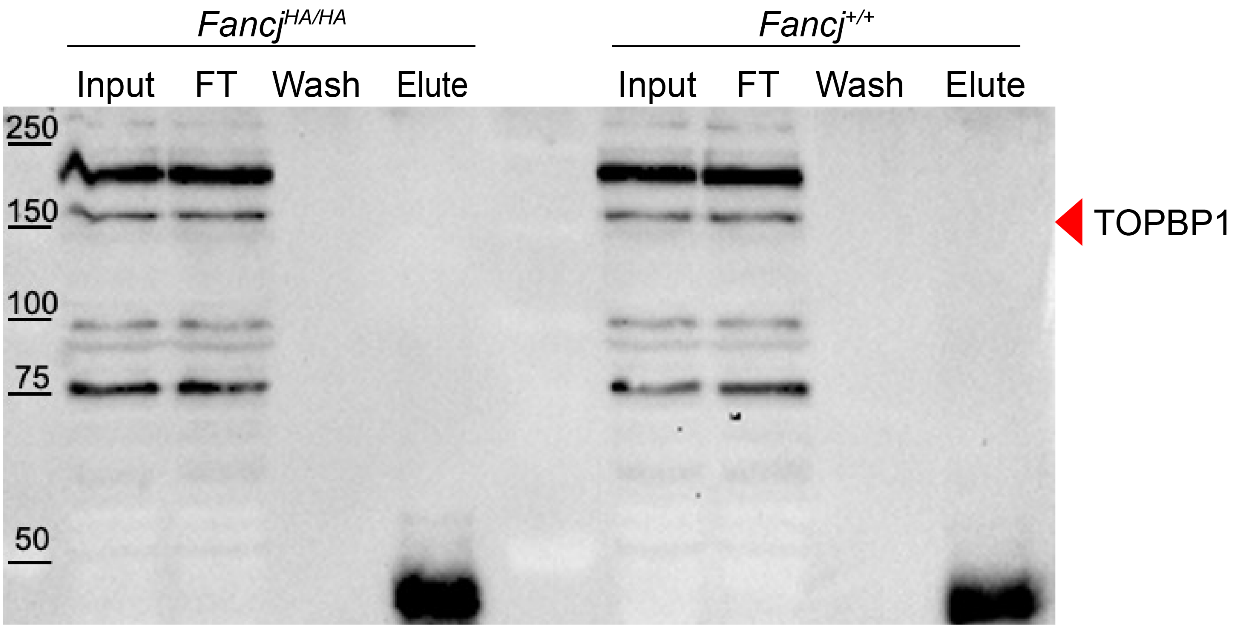


**Figure S5 FANCJ does not interact with TOPBP1 in meiosis.** Western blot of HA-IP using whole testis lysate from *Fancj*$^{HA/HA}$ and *Fancj*$^{+/+}$mice. Membrane was blotted with antibody against TOPBP1. Red arrow points to expected TOPBP1 band size (~169 kDa).
